# Genetic resistance to DEHP-induced transgenerational endocrine disruption

**DOI:** 10.1101/474155

**Authors:** Ludwig Stenz, Rita Rahban, Julien Prados, Serge Nef, Ariane Paoloni-Giacobino

## Abstract

Di(2-ethylhexyl)phthalate (DEHP) interferes with sex hormones signaling pathways (SHP). C57BL/6J mice prenatally exposed to DEHP develop a testicular dysgenesis syndrome (TDS) at adulthood, but similarly-exposed FVB/N mice are not affected. Here we aim to understand the reasons behind this drastic difference that should depend on the genome of the strain. In both backgrounds, pregnant female mice received *per os* either DEHP or corn oil vehicle and the male filiations were examined. Computer-assisted sperm analysis showed a DEHP-induced decreased sperm count and velocities in C57BL/6J. Sperm RNA sequencing experiments resulted in the identification of the 62 most differentially expressed RNAs. These RNAs, mainly regulated by hormones, produced strain-specific transcriptional responses to prenatal exposure to DEHP; a pool of RNAs was increased in FVB, another pool of RNAs was decreased in C57BL/6J. In FVB/N, analysis of non-synonymous SNP impacting SHP identified rs387782768 and rs387782768 respectively associated with absence of the Forkhead Box A3 (*Foxa3*) RNA and increased expression of estrogen receptor 1 variant 4 (NM_001302533) RNA. Analysis of the role of SNPs modifying SHP binding sites in function of strain-specific responses to DEHP revealed a DEHP-resistance allele in FVB/N containing an additional FOXA1-3 binding site at rs30973633 and four DEHP-induced beta-defensins (*Defb42*, *Defb30*, *Defb47* and *Defb48*). A DEHP-susceptibility allele in C57BL/6J contained five SNPs (rs28279710, rs32977910, rs46648903, rs46677594 and rs48287999) affecting SHP and six genes (*Svs2*, *Svs3b*, *Svs4*, *Svs3a*, *Svs6* and *Svs5)* epigenetically silenced by DEHP. Finally, targeted experiments confirmed increased methylation in the *Svs3ab* promoter with decreased SEMG2 persisting across generations, providing a molecular explanation for the transgenerational sperm velocity decrease found in C57BL/6J after DEHP exposure. We conclude that the existence of SNP-dependent mechanisms in inbred mice may confer resistance to transgenerational endocrine disruption.

## Introduction

Genetic factors combined with epigenetic mechanisms disrupting androgen actions during fetal life may result in testicular dysgenesis syndrome (TDS) at adulthood. TDS is characterized by poor semen quality, urogenital malformations and testicular cancers [1]. Phthalate plasticizers are well-described detrimental environmental factors interfering with androgen actions and considered as repro-toxic. Both di-(2-ethylhexyl) phthalate (DEHP; CAS No. 117-81-7) and its principal metabolite, mono- (2-ethylhexyl) phthalate (MEHP; CAS No. 4376-20-9) interact with the androgen (AR), estrogen (ER) and peroxisome proliferator-activated receptors (PPARs), with a negative impact on testosterone production [2, 3]. Multiple direct effects at the molecular level of both DEHP and its metabolite were observed *in vitro* by using a reporter assay with human cell lines on the estrogen receptor (ESR1) and on AR [2]. Accumulated data thus demonstrate that DEHP interferes with sex steroid hormone signaling pathways (SHP).

The impact of prenatal exposure to DEHP on human reproductive health at adulthood remains largely unknown as human prospective studies testing such associations are lacking. However, an increasing body of evidence shows that prenatal exposure to DEHP may cause androgen deficiency during embryogenesis, with effects in animal models reproducing those observed in humans. First, a direct negative impact of DEHP on testosterone production was reported in the rat fetal testis after prenatal exposure to DEHP, as well as in human testis explants cultured with DEHP or MEHP [4, 5]. Second, meta-analyses in human studies resulted in a statistically significant association between maternal urinary concentrations of DEHP metabolites and a shortened anogenital distance (AGD) in boys, consistent with the statistically significant association obtained between DEHP treatment and reduced AGD in animal meta-analysis [6]. Currently, AGD remains the best marker of fetal androgen exposure in humans [7]. DEHP and MEHP concentrations in maternal urine during pregnancy, as well as in umbilical cords, were associated with a reduced AGD in male newborns, but also with a shortened gestation period [8, 9].

Interestingly, a dose-response gradient and heterogeneity explained by strains were observed in animal studies [6]. Our previous studies in mice suggest that genetic variations may explain a strain-dependent alteration in adult sperm induced by prenatal exposure to DEHP, probably causing an endocrine disruption during the development of the embryo. We administered DEHP to pregnant C57BL/6J and FVB/N inbred mice strains to evaluate the impact of prenatal exposure to DEHP on sperm counts, as well as on the epigenetic status of the male gametes produced at adulthood by the first filial generation of exposed animals (F1) [10]. DEHP was injected *per os* to pregnant mice at the dose of 300 mg/kg/day and during embryonic (E) days (E9-19). The dose was chosen from a previous study and appears to be relevant for extreme human exposure, corresponding to 16 mg/kg/day of direct exposure in preterm infants treated in neonatal intensive care units, when taking into account all maternal barriers existing between the injection site and the embryos in the mice model [11]. The exposure time frame extends from the primordial germ cell (PGC) migration period (~E10.5) and covers the gonadal differentiation period initiated between E11 and E12 in the developing mice embryos, representing a susceptible time window for androgen interference. In our previous study, a decreased sperm count was observed in the C57BL/6J strain, but not in FVB/N mice, indicating that the latter seem to be resistant and the former sensitive to DEHP [10]. Thus, we postulated that genomic variations should be responsible for strain-specific impacts.

Both DNA methylation and RNA expression may be affected in the sperm of mice prenatally exposed to DEHP. We showed that PGCs were present at the time of DEHP exposure. PGCs are characterized by a specific pattern of DNA methylation and transcription controlling the cell fate in the lineage and involving notably genes required for pluripotency, as well as germline-specific genes. PGCs differentiate themselves in male germ cells during the fetal androgen-dependent sexual differentiation of the somatic gonads to testis. At puberty, a peak of testosterone triggers the spermatogenesis process starting from the spermatogonia. The latter is derived from the male fetal germ cell by mechanisms still poorly understood. The spermatogenesis involves transcriptional patterns specific to the successive developmental stages from spermatogonia to spermatozoa [12]. Spermatogonia undergo mitotic divisions with incomplete cytokinesis in the testis, producing spermatids in the lumen of the seminiferous tubule, then maturing in spermatozoa in the epididymis. Mature spermatozoa located in the cauda epididymis are those with the ability to fertilize an egg naturally after ejaculation. In mature sperm, RNAs remain present despite inactive transcription, with some deeply embedded in the sperm nucleus. At least part of these paternal RNAs is transmitted to the egg at fertilization, whereas DNA methylation changes occur during the entire spermatogenesis process. Thus, DNA methylation and RNA expression appear to be dynamically controlled across the entire “lifetime” of the male germ cell lineage. Under such circumstances, endocrine disruption taking place during the susceptible time frames of PGC migration may probably alter DNA methylation and RNA expression status in mature sperm produced at adulthood in a long-term manner.

Here, we first postulate that FVB/N resistance or alternatively C57BL/6J susceptibility, may involve genomic variations affecting the androgen signaling occurring in a strain-specific manner in the sperm. To test this hypothesis, comparison between both genomes performed by Wong K. *et al* were incorporated in the present study [13]. In mature spermatozoa, transcription is arrested and DNA methylation is lower than in differentiated cells, but a functional AR is expressed in both X and Y carrier spermatozoa, dihydrotestosterone (DHT) is present in the seminal fluid, and steroid receptors and their ligands may impact on male gamete functions [14, 15]. Results revealed that the AR is not directly affected by polymorphisms in FVB/N compared with the C57BL/6J reference genome. However, the non-synonymous single nucleotide polymorphism (SNP) rs29315913 was found in one variant of ESR1 and rs387782768 was found in the Forkhead box A3 (FOXA3) transcription factor required for testicular steroidogenesis [13, 16]. Briefly, ESR1 is involved in the regulation of male reproductive organs, notably the efferent ductus and the epididymis, and its loss results in impaired ion transport and water reabsorption, with production of abnormal sperm characterized by abnormal flagellar coiling and increased incidence of spontaneous acrosome reactions [17, 18]. The Forkhead box A genes (FOXA1, FOXA2 and FOXA3) encode pioneer transcription factors, i.e. the first detectable factors to engage target sites in chromatin, thus facilitating the further binding of the AR in the nucleus to the targeted recognized DNA sequences [19].

The aim of this genome-environment interaction study focalized on male germ cells was to identify the SNPs that may be responsible for changes in sperm RNA content, which depend on the strain and on DEHP exposure, among those varying between FVB/N and C57BL/6J inbred mice as it is already known that some slight differences exist between both strains in the sexual hormone receptors and their FOXA-associated factors. We investigate also the transmission of biological alterations to subsequent generations by altered sperm, as well as the epigenetic status of genes encoding the dysregulated RNAs.

## Results

### Phenotypic impact of *in utero* exposure to DEHP in C57BL/6J and FVB/N mice

The phenotypic changes affecting male fertility parameters induced by prenatal exposure to DEHP confirm the FVB/N resistance found in our earlier study using an independent method [10] and using data partially previously obtained in C57BL/6J [20]. In FVB/N, AGD, testes weight, sperm concentration and curvilinear velocity (VSL) were not affected by prenatal exposure to DEHP and remain stable also in the second generations of mice generated from the prenatally exposed F1 males (Fig 1AB). The only exception concerns sperm velocities with decreased average path velocity (VAP) and VSL in FVB.D300.F1 compared to FVB.CTL.F1. A return to control values was observed in FVB.D300.F2 (Fig 1B). The experiments were stopped at F2 in FVB/N due to the absence of any remaining impact on all tested parameters.

**Fig 1.**
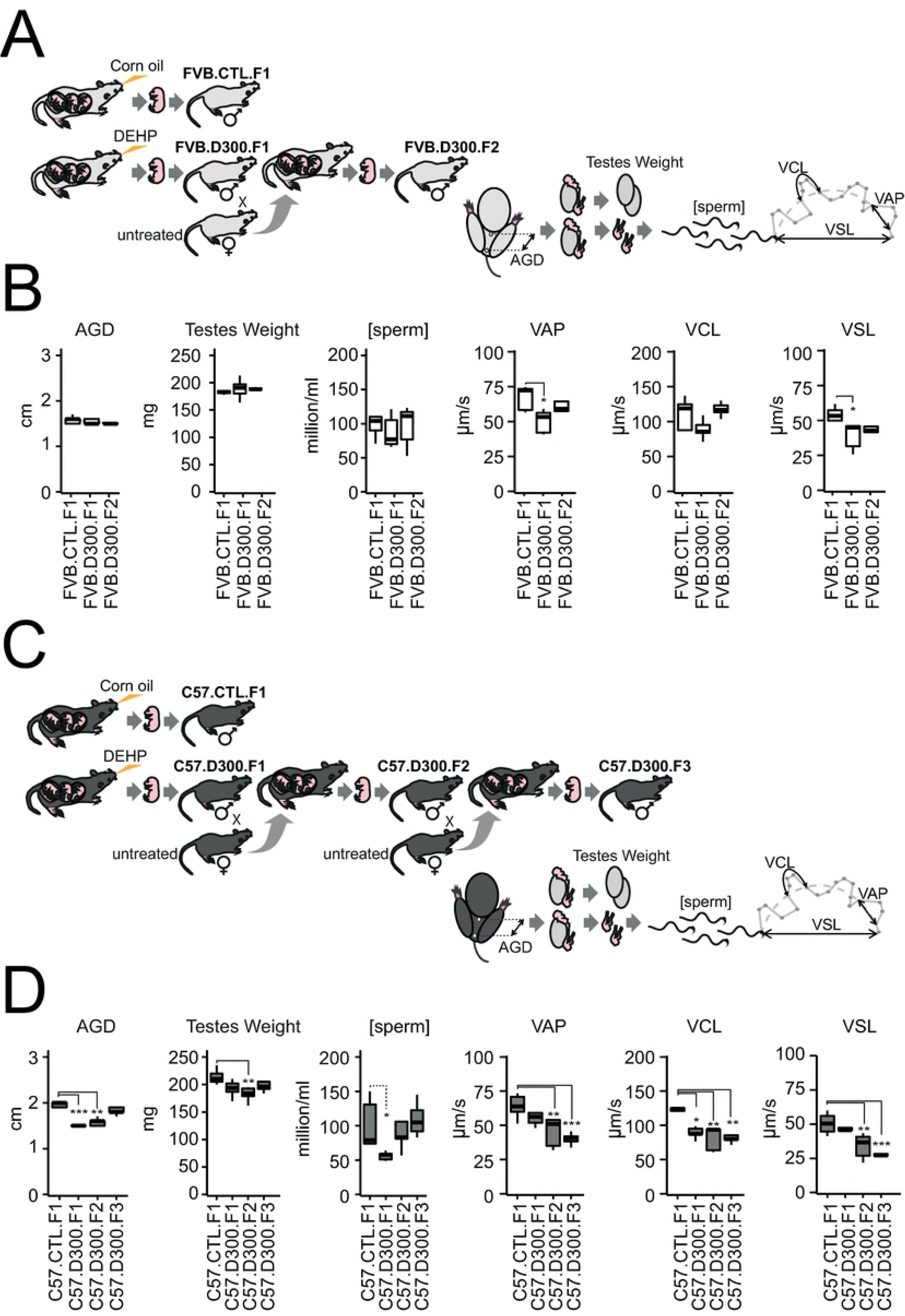
Strain-dependent alterations of fertility parameters in male offspring exposed in utero to DEHP. *Prenatal exposures to DEHP were performed daily from E9-19 in both FVB/N and C57BL/6J mice strains using per os injections in hand-restrained pregnant females. Injections of 20 µl of 1.15 M DEHP diluted in corn oil corresponding to 300 mg/kg/day (D300) for 30 g mice estimates were compared with injections of 20 µl of corn oil as a vehicle for a fat-soluble compound, settled as the control condition (CTL). Male fertility parameters were obtained at P100 using five biological replicates per condition. AGD and testes weight were obtained prior to computer-assisted sperm analysis (CASA). CASA was used to assess sperm concentrations expressed in million per ml [sperm] and sperm velocities expressed in µm per second (µm/s). VAP: average path velocity; VCL: curvilinear velocity; VSL: straight-line velocity. (A)* Experimental design of the study performed in the strain FVB/N. *FVB.CTL.F1 were born from the corn oil-exposed mother. FVB.D300.F1 males were born from the DEHP-exposed mother. FVB.D300.F2 males were obtained after mating with unexposed females with FVB.D300.F1.* (B). Male fertility parameters obtained in FVB after prenatal exposure to DEHP. *AGD, testes weight, sperm concentrations and VCL did not differ across the tested conditions. VAP (p-value <0.02) and VSL (p<0.02) were decreased in FVB.D300.F1 compared with FVB.CTL.F1.* (C). Experimental design of the study performed in the strain C57BL/6J. *C57.CTL.F1 males were born from the corn oil-exposed mother, whereas C57.D300.F1 males were born from the DEHP-exposed mother. C57.D300.F2 males were obtained after mating with an unexposed female with C57.D300.F1. C57.D300.F3 males were obtained after mating with an unexposed female with C57.D300.F2 males.* (D) Male fertility parameters obtained in C57BL/6J strain after prenatal exposure to DEHP. *Compared with C57.CTL.F1, AGD, sperm concentration and VCL were decreased in C57.D300.F1 (p <7*10*^−5^, *p <0.05, p <0.02, respectively), AGD (p < 0.002), testes weight (p <0.008), VAP (p <0.008), VSL (p <0.008) and VCL (p <0.003) were decreased in C57.D300.F2, and VAP (p <1*10*^−3^), *VSL (p <1*10*^−3^) *and VSL (p<0.004) were decreased in C57.D300.F3. Statistical significance was performed using a one-way analysis of variance with a post-hoc Tukey honestly significant difference test (ANOVA-Tukey HSD). Significant differences between the tested conditions and the control conditions are reported as a graphic. * corrected p-value <0.05, ** corrected p-value <0.01, *** corrected p-values <0.001*.

In contrast, all parameters tested in C57BL/6J were affected by prenatal exposure to DEHP in both the first and second generations in the DEHP-exposed lineage (Fig 1CD). These results are compatible with a DEHP-induced TDS. AGD, sperm concentration and VCL were decreased in C57.D300.F1, and AGD, testes weight, VAP, VCL and VSL were decreased in C57.D300.F2. These multigenerational detrimental impacts of prenatal exposure to DEHP affecting various male fertility parameters are compatible with an intergenerational inheritance in C57BL/6J. Surprisingly, a continuous deterioration of sperm velocity was observed in the DEHP-exposed lineage in the C57BL/6J strain, compatible with a transgenerational increased inheritance of decreased sperm velocity (Fig 1D). At F3 in C57BL/6J, VAP decreased by 24 µm/s between C57.F1.CTL and C57.F3.D300 (p-value = 8.8*10^−4^), VCL decreased by 36 µm/s between F1.CTL and F3.D300 (p-value = 3.6*10^−3^), and VSL decreased by 24 µm/s between F1.CTL and F3.D300 (p-value = 2.4*10^−4^).

Overall, the results demonstrated a strain-specific difference in the impact of prenatal exposure to DEHP in inbred mice. They also strongly suggest the persistence of a detrimental impact across generations in the case of DEHP-susceptibility and revealed a surprising deterioration of sperm velocity apparently inherited in a transgenerational manner.

### Sperm transcriptome variations induced by prenatal exposure to DEHP in C57BL/6J and FVB/N mice

SHP were shown to be affected by DEHP and its metabolites in previous reports. Both androgens and estrogens exerted their biological effects on gene transcription upon binding to their respective receptors (AR and ESR1). Here, we postulate that prenatal exposure to DEHP may dysregulate sperm RNAs in a persistent manner. Total RNA was extracted from sperm samples analyzed by CASA and the RNA content was analyzed using all-RNA-seq (Fig 2A and Table S1). One differential analysis was performed using multiple pairwise comparisons between the conditions C57.CTL.F1, C57.D300.F1, FVB.CTL.F1 and FVB.D300.F1 in order to take into account the impact of the strains on sperm RNA content in the absence or presence of DEHP exposure. Additionally, as almost all parameters tested remained affected at F2 in C57BL/6J (Fig 1), we also included the C57.D300.F2 condition.

**Fig 2.**
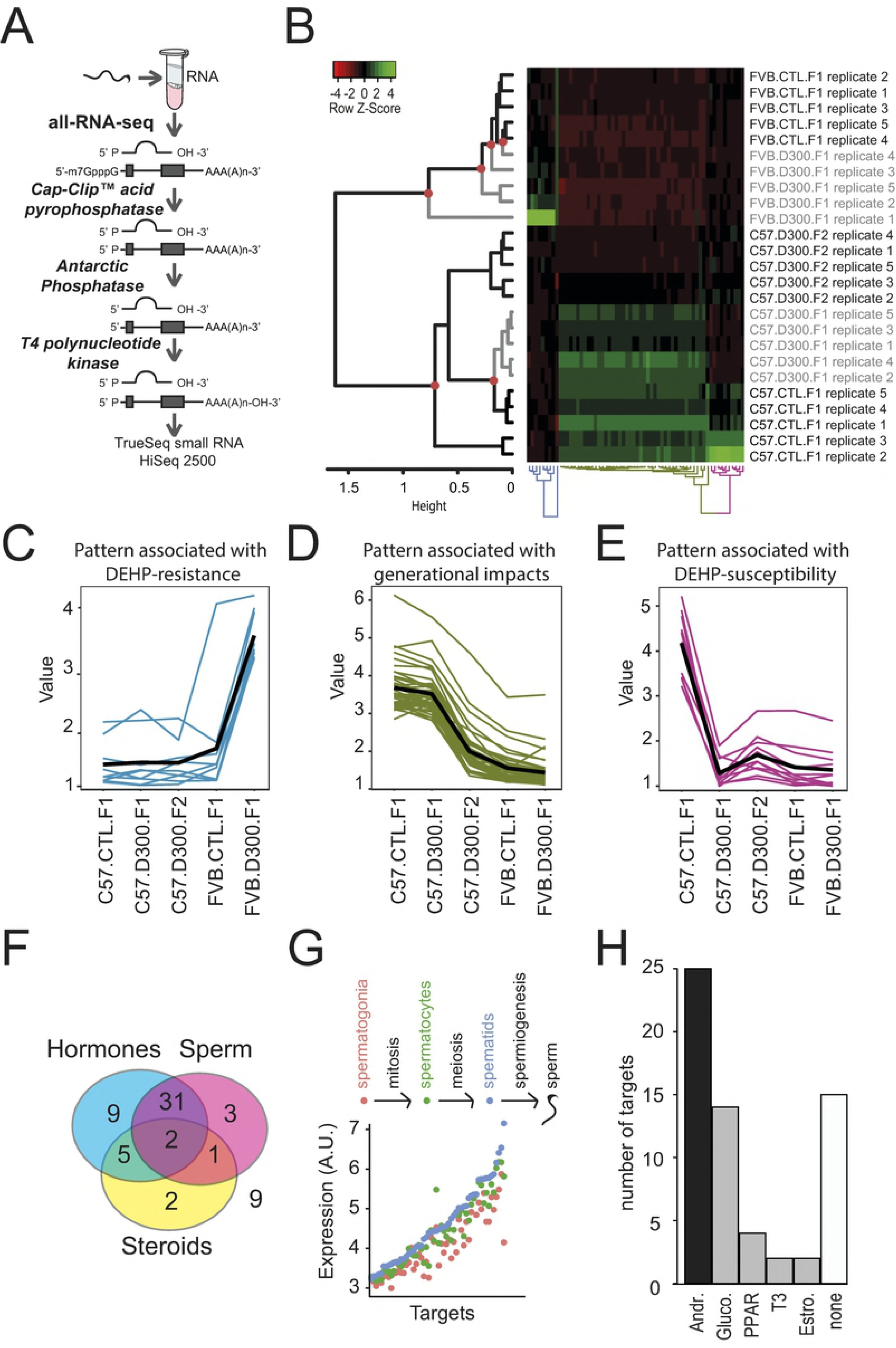
Results obtained in the genome*environment*generation transcriptomic interactions study. (A) All-RNA-seq analysis in sperm. Sperm sampled from the cauda epididymis was analyzed in quintuplicates using the all-RNA-seq protocol, which allows measurements of all RNA types independently of their end variations. (B) Hierarchical clustering analysis combined with a heat-map illustration of all-RNA-seq data. Red circles: significant nodes in the dendrogram. In the heat-map, high levels of expression are in green and low levels in red. (C) The expression pattern associated with the DEHP-resistance indicates induction of expressions in FVB.D300.F1 compared with FVB.CTL.F1. (D) Pattern associated with generational impacts in the C57BL/6J background. Decreased expressions are observed in C57.D300.F2 compared with C57.CTL.F1 and C57.D300.F1. (E) The expression pattern associated with DEHP-susceptibility displayed decreased expressions in C57.D300.F1 compared with C57.CTL.F1. (F) Triple-Venn diagram segregating the 62 RNAs. The 62 RNAs were segregated in a Triple-Venn diagram as hormonally-regulated (light blue circle: N=47; 76%), as previously identified in sperm (pink circle: N=37; 60%), or involved in the transport of cholesterol or the metabolism of steroids (yellow circle: N=10; 16%). The majority of RNAs are hormonally-regulated and expressed in sperm (N=33; 53%); the minority of RNAs are out of the diagram (N=9; 15%). (G) Expression levels of RNA across germ cells subtypes: spermatogonia in red, spermatocytes in green, and spermatids in blue. RNAs were significantly enriched for increased expression in spermatogenesis compared with all RNAs in the Chalmel dataset (χ²(1) = 53; p< 3.3*10^−13^). (H) Number of RNA regulated by androgen (Andr.), glucocorticoids (Gluco.), hormone-like PPARs, triiodothyronine (T3), estrogen (Estro.) or not regulated by hormones (none). 76% of the targets present a reported hormonal regulation (47/62), among these 53% by androgen (25/47).

This approach resulted in genome*environment*generation interactions transcriptomic analysis identifying 62 RNAs as the most differentially expressed RNA among the different conditions (Fig 2B). Importantly, three different patterns of expression changes were found (Figs 2C, 2D and 2E). The first pattern showed increased RNA levels specifically in FVB.D300.F1 compared with FVB.CTL.F1. This pattern presenting an FVB/N-specific and DEHP-mediated induction of RNA expression was associated with FVB/N resistance (Fig 2C). The second pattern was associated with generational changes occurring between the first and second generations in the C57BL/6J exposed lineage (Fig 2D). The third pattern revealed decreased levels of various RNA specifically in C57.D300.F1 compared with C57.CTL.F1, without changes in FVB/N. The latter pattern was associated with C57BL/6J susceptibility to DEHP (Fig 2E). Overall, the 62 differentially expressed RNA across the tested conditions were mainly regulated by hormones in sperm or in male reproductive tissues (Fig 2F and Table 1). The vast majority (80%) present increased expression during spermatogenesis (Fig 2G); androgen was the most represented regulatory hormone (Fig 2H). In addition, pathways highly relevant to sperm physiological functions and DEHP toxicity were affected across the tested conditions (Table S2).

**Table 1.**
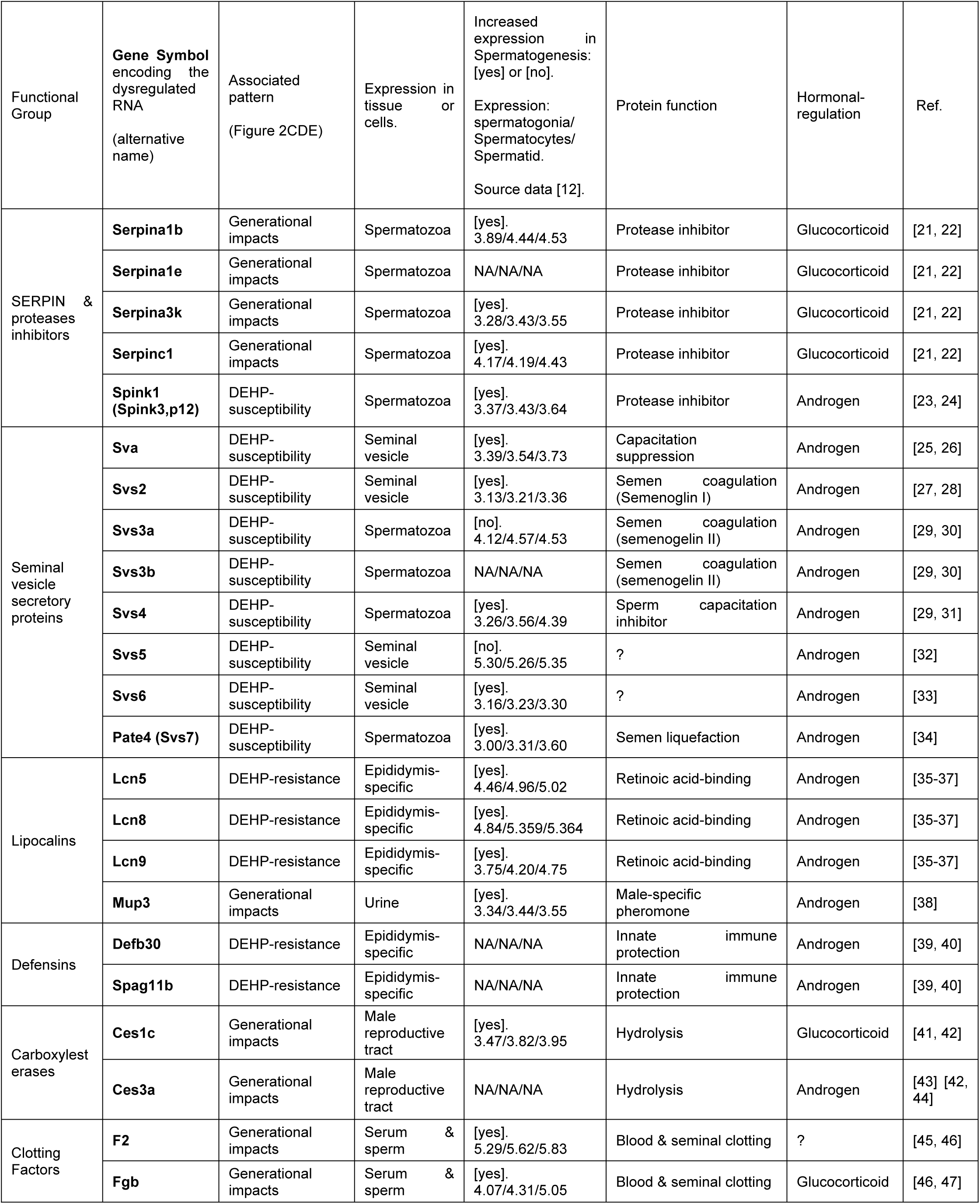
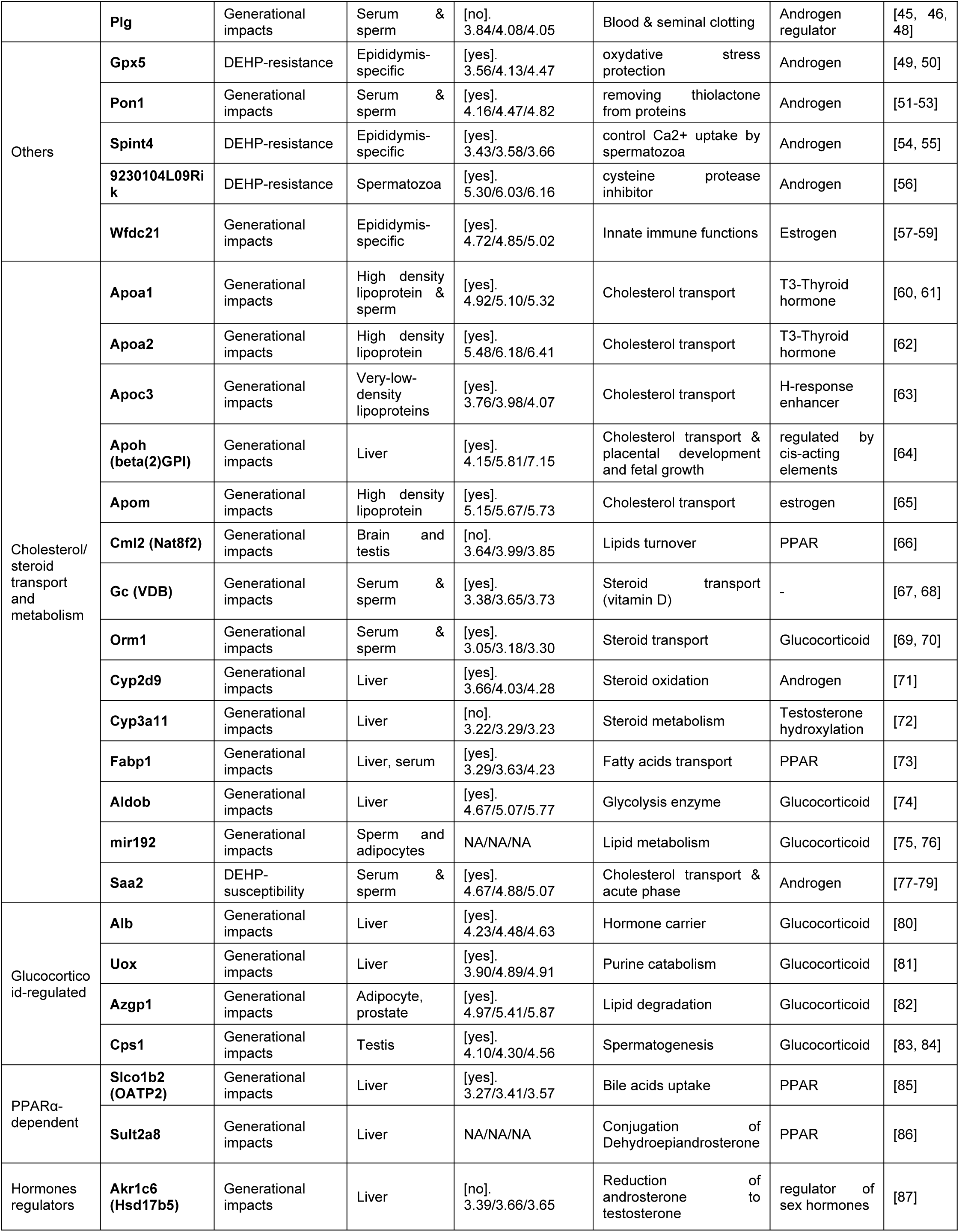
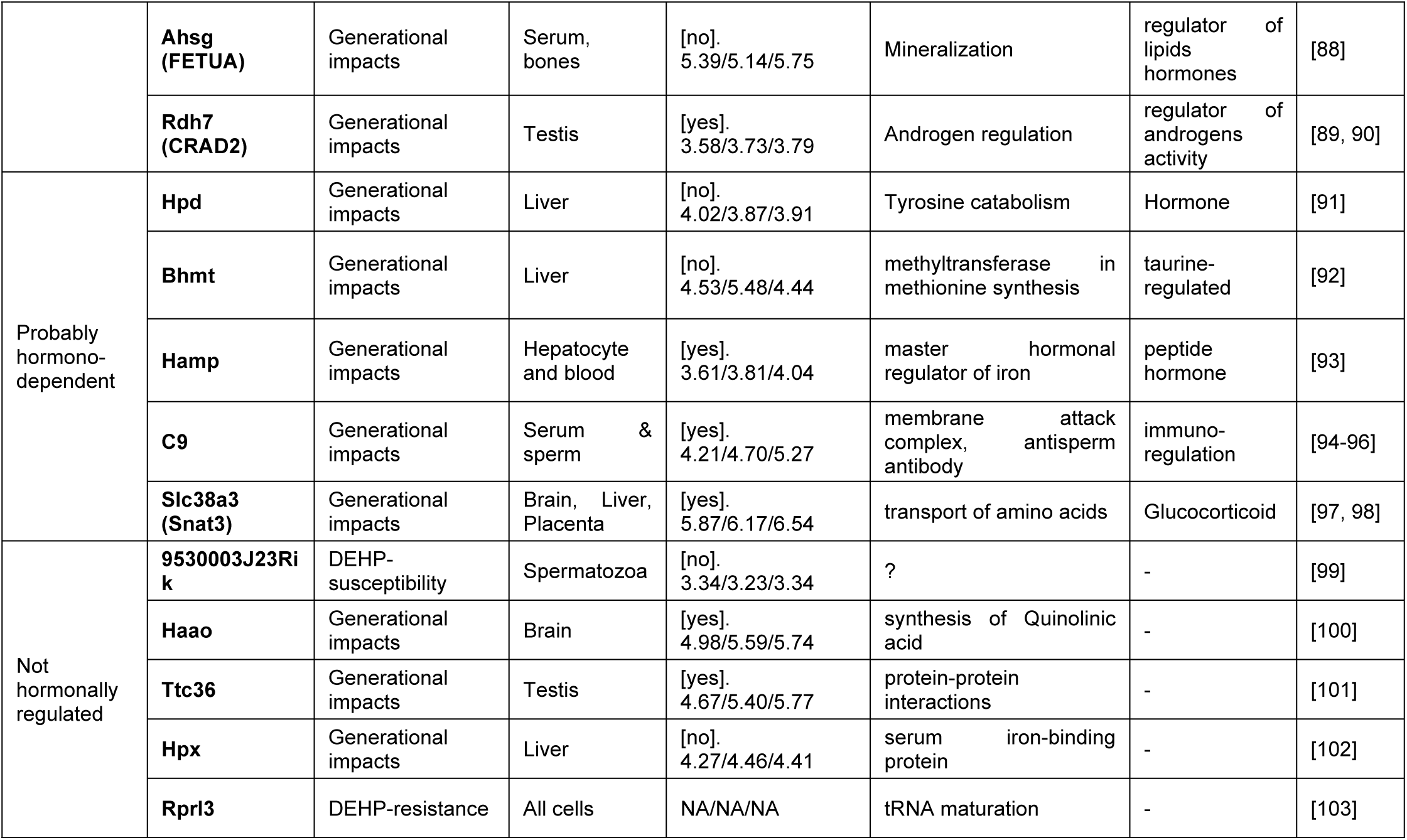
Characterization of the 62 dysregulated RNAs.

Overall, these results demonstrate that prenatal exposure to DEHP may alter the male meiotic program in a strain-specific manner, producing distinct patterns of expression changes and mainly implying dysregulation in androgen-regulated sperm RNAs.

### Strain-specific non-synonymous SNPs affecting AR, ESR1 and FOXA1-3 sperm RNA levels

SNP variations between FVB/N and C57BL/6J affecting SHP may explain the strain-specificity of the DEHP-induced alterations in sperm RNAs shown in Fig 2. Non-synonymous SNP rs387782768 in FOXA3 and rs29315913 in ESR1 variant 4 genes were characterized previously [13]. No other relevant SNP affecting SHP so directly could be found in both strains. Transcriptomic results of this study revealed that both SNPs were associated with strain-specific differences of particular RNA expression levels, independent of DEHP exposure (Fig 3A). First, rs387782768 was associated with the absence of FOXA3 sperm transcripts in FVB/N. Second, rs29315913 was associated with increased levels of ESR1 variant 4 sperm transcript in FVB/N. Other tested transcription factors not affected by non-synonymous SNP (FOXA1, FOXA2, AR and other ESR1 variants) did not differ in their expression levels between both strains. An upregulation of AR expression was also observed in C57.D300.F2 that did not depend directly on a SNP, apparently reflecting a generational impact in the exposed lineage of the C57BL/6J strain.

**Fig 3.**
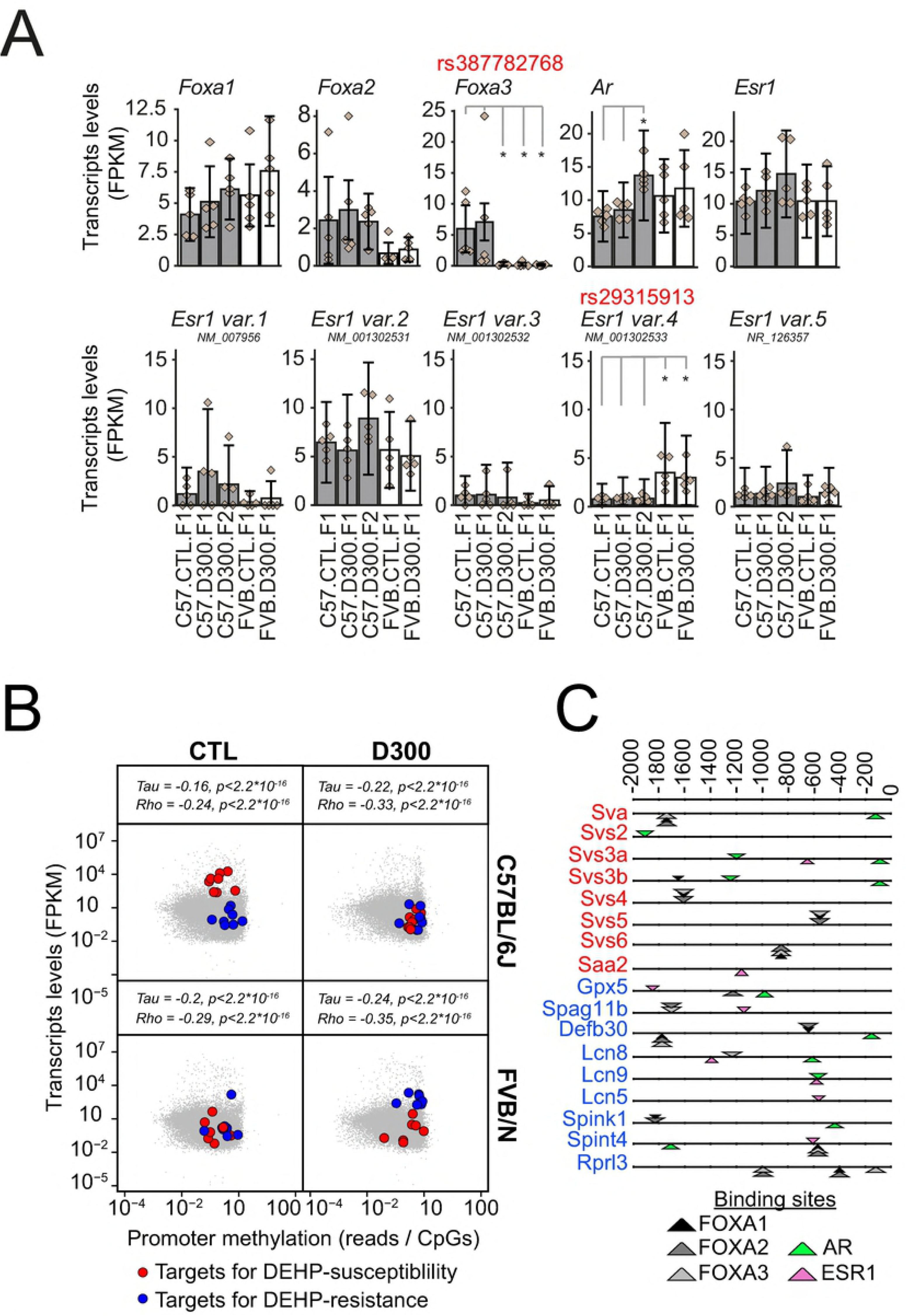
Sexual hormone signaling differs between FVB/N-resistant and C57BL/6J-susceptible strains. (A) Impact of non-synonymous SNP on SHP key players. Expression levels for FOXA1-3 transcription factors, as well as for AR and ESR1, across the five experimental conditions. C57BL/6J-derived data are shown as gray bars and FVB/N derived data as white bars. Error bars represent the CummeRbund computed confidence intervals. Diamonds show the individual FPKM values of the replicates. Data were produced by all RNA-seq. Non-synonymous rs387782768 in FOXA3 and rs29315913 SNP in ESR1 variant 4 were characterized previously [13]. Stars represent statistically significant differences of expression levels according to the non-parametric pairwise Wilcoxon test with Benjamini & Hochberg correction for multiple testing. (B) Promoter methylation and RNA expression in the sperm of the F1 generation of C57BL/6J (up) and FVB/N (down) mice subjected to prenatal exposure to corn oil vehicle (CTL, left) or to DEHP (D300, right) were analyzed in quintuplicates. Promoter methylations were obtained by MBD-seq [10] and were normalized to the number of CpG sites in 2,2 Kb. Correlations were assessed using Kendall’s tau and Spearman’s rho rank correlation tests. Blue points represent the genes showing the pattern of DEHP-resistance involving Gpx5, Spag11b, Defb30, Lcn8, Lcn9, Lcn5, Spint4 and Rprl3, that was absent in the MBD-seq, but without 9230104L09Rik. Red points represent the genes showing the pattern of DEHP-susceptibility involving Sva, Svs2, Svs3a, Svs3b, Svs4, Svs5, Svs6, Spink1 and Saa2, without Pate4 and 950003J23Rik, both absent in MBD-seq. (C) Binding sites for AR (green), ER (pink) and FOXA1-3 transcription factors (black, dark gray and white) detected in promoters of genes associated with DEHP-susceptibility (red names) and DEHP-resistance (blue names). The number of AR (n=6) binding sites were higher than ER (n=2) sites in promoters of genes (n=8) silenced by DEHP in C57BL/6J and associated with susceptibility. Equal numbers of ER (n=6) and AR (n=6) binding sites were detected in the genes (n=9) induced by DEHP in FVB/N and associated with resistance. Binding sites are shown as triangles orientated differently depending on the DNA strands and with thickness reflecting the score. Sequences were extracted from −2000 to 0 bp with gene names from Mus musculus GRCm38. Analysis was performed using the matrix-scan program of Rsat [105].

These results revealed that strain-specific SNP variations affecting key players in the SHP resulted in differences in sperm RNA levels and they may play a role in the observed strain-specific susceptibility to hormonal disruption.

### Effects of DEHP on promoter methylation and on gene expression in the sperm of both strains

Changes in RNA levels observed in sperm upon prenatal exposure to DEHP may involve alteration in epigenetic mechanisms. Among these, aberrant DNA methylation(s) in the male germ cell lineage caused by prenatal DEHP exposure may result in persistent alterations of sperm RNA patterns detectable at adulthood. To test this hypothesis, all-RNA-seq data were merged to previously acquired Methyl-CpG-binding domain sequencing (MBD-seq) data for all conditions tested at F1 also performed in quintuplicates (C57.CTL.F1, C57.D300.F1, FVB.CTL.F1 and FVB.D300.F1) [10]. The first observation is that transcript abundance was negatively correlated with promoter methylation in the sperm, consistent with a repressive impact of promoter-methylation on transcription named silencing (Fig 3B) [106]. Promoter methylation was then tested specifically in the RNA that produced both patterns associated with either DEHP-susceptibility or DEHP resistance (Figs 2C and 2E, respectively, and Table 1). The targets for DEHP-susceptibility, characterized by DEHP-induced decreased RNA expression levels, were all associated with a DEHP-induced increased promoter methylation in C57BL/6J (Fig 3B, red points). These combined results are consistent with a DEHP-induced epigenetic silencing of the DEHP-susceptible targets identified only in C57BL/6J, as the same targets were not affected by DEHP in FVB/N, neither in their expression, nor in their promoter methylations levels (Fig 3B, red points). The targets for DEHP-resistance, upregulated by DEHP specifically in FVB/N, were also associated, paradoxically, with promoter methylation increases (Fig 3B, blue points). The targets for DEHP-resistance were not affected by DEHP in C57BL/6J, neither in their expressions, nor in their promoter methylation levels (Fig 3B, blue points).

We postulated that the decreased expression of androgen-regulated genes in particular may be directly explained by the anti-androgenic activity of DEHP in C57BL/6J, whereas the increased expression of genes in FVB/N may involve an additional pro-estrogenic activity of DEHP. Therefore, the binding sites for both estrogen and androgen signaling systems were analyzed in the promoters of the identified targets for both DEHP-susceptibility and DEHP-resistance (Fig 3C). Results revealed that receptor binding sites for androgen signaling were higher than for estrogen signaling in the promoters of targets for DEHP-susceptibility (Fig 3C, red gene symbols), whereas equal numbers of ER and AR binding sites were detected in the promoters of targets for DEHP-resistance (Fig 3C, blue gene symbols). These results are compatible with anti-androgenic silencing and pro-estrogenic induction of the strain-specific targets.

Overall, these results demonstrate increased promoter methylations in strain-specific affected targets for DEHP-susceptibility in C57BL/6J, and for DEHP-resistance in FVB/N, with differences in the number of ER over AR binding sites found in the promoters of these strain-specific targets.

### Impact of SNPs on strain-specific DEHP-induced sperm RNA changes

SNP variations affecting binding sites for FOXA1-3, AR and ESR1 and transcription factors were investigated in function of the DEHP-mediated, strain-specific sperm RNA changes (Fig 4). Such an approach may allow the identification of alleles associated with either DEHP-susceptibility or DEHP-resistance if several SNPs are co-localized or if various co-localized genes are affected.

**Fig 4.**
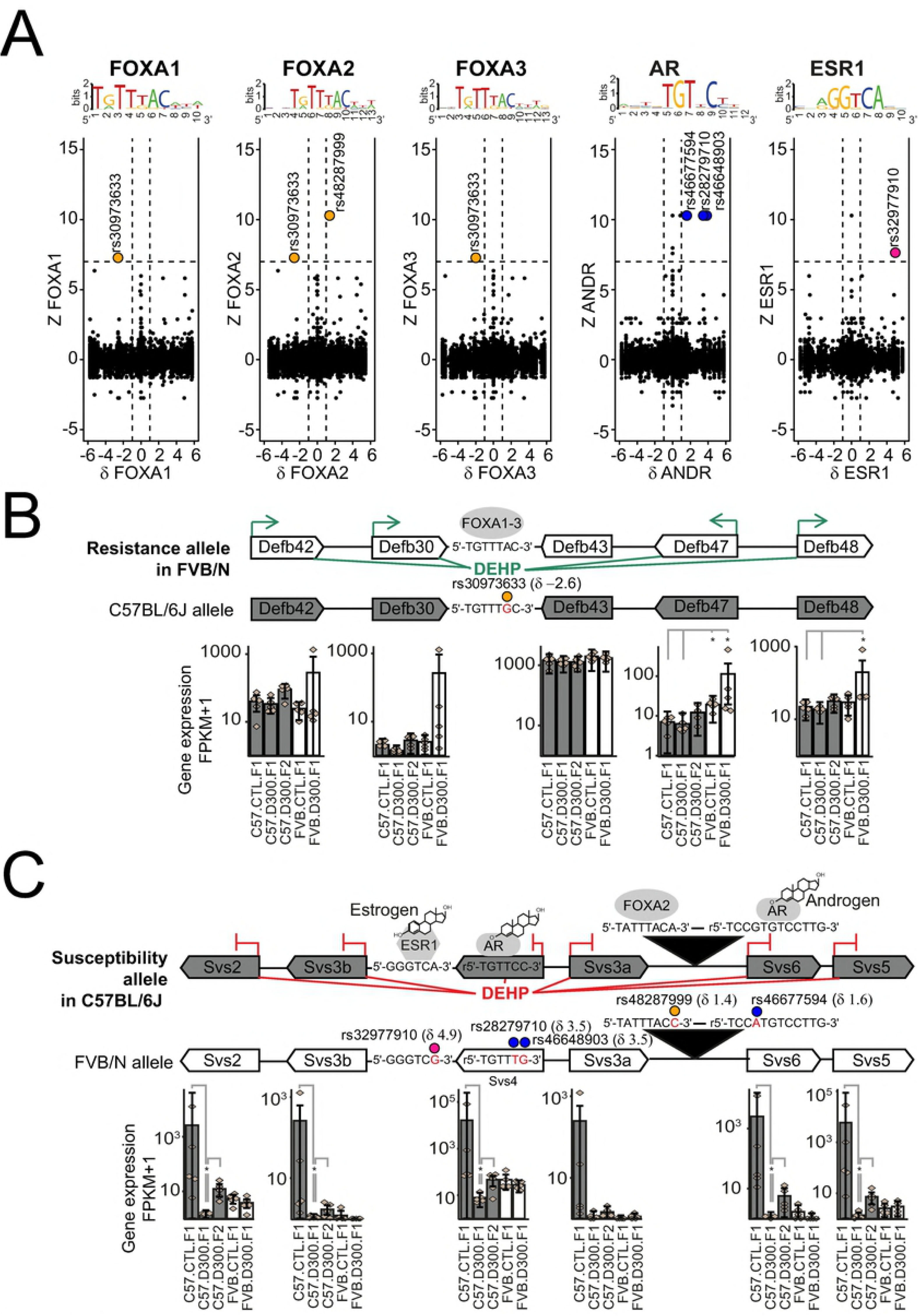
Genome-wide analysis of SNP impacting HRP motifs in function of strain-specific dysregulation of male germ cell RNAs. (A) Genome-wide analysis of DEHP-induced and strain-specific transcriptional changes (Z) in the function of strain-specific binding site (δ) for FOXA1, FOXA2, FOXA3, ANDR and ESR1. Z = log2 (FPKM _FVB.D300.F1_ / FPKM _FVB.CTL.F1_) - log2 (FPKM _C57.D300.F1_/ FPKM _C57.CTL.F1_). δ = score _C57BL/6J_ –score _FVB/N_. The corresponding logos above each graph represent the tested motif. Dashed lines represent thresholds. (B) FVB/N-specific DEHP-induced increased expression in the β-defensin loci associated with an FVB/N-specific FOXA1-3 binding motif due to rs30973633. Upon prenatal exposure to DEHP, increased expression levels are recorded for Defb42, Defb30, Defb47 and Defb48, but not for Defb43 in FVB/N, without changes in C57BL/6J. (C) C57BL/6J-specifc DEHP-induced expressional changes in the Svs loci associated with C57BL/6J-specific binding sites for AR, ESR1 and FOXA2 due to SNPs rs28279710, rs46648903, rs46677594, rs32977910 and rs46677594. In utero exposure to DEHP resulted in decreased expression across all Svs genes in C57BL/6J with no change in FVB/N. (BC) Delta values in parentheses represent score differences between both alleles for the respective motif, taking the SNP into account. Stars represent statistically significant differences of expression levels, according to the non-parametric pairwise Wilcoxon test with Benjamini & Hochberg correction for multiple testing. In part C, statistically significant differences between both strains are not shown.

A total of 6 SNPs showed the highest combined impacts on both the strain-specific transcriptional changes induced by prenatal exposure to DEHP measured with Z values, as well as on HSP-motifs specificities measured with δ values (Fig 4A).

We were able to identify rs30973633 as affecting a binding motif for all FOXA1-3 transcription factors in that this motif is present in the FVB/N and absent in the C57BL/6J allele. Moreover, the FVB/N-specific binding motif is associated via a probable DEHP-mediated induction of FOXA1-3 (Fig 3A), with the activation of various beta-defensins localized around rs30973633 (Fig 4B and Table 2). Precisely, prenatal exposure to DEHP induced increased expressions in *Defb42*, *Defb30*, *Defb47* and *Defb48*, but not *Defb43*. None of these genes were affected in their expression after prenatal exposure to DEHP in C57BL/6J (Fig 4B). This DNA region displaying an FVB/N specific induction of RNA expression combined with an FVB/N specific binding site for FOXA1-3 was defined as an “FVB/N resistance allele” to prenatal DEHP exposure.

**Table 2.**
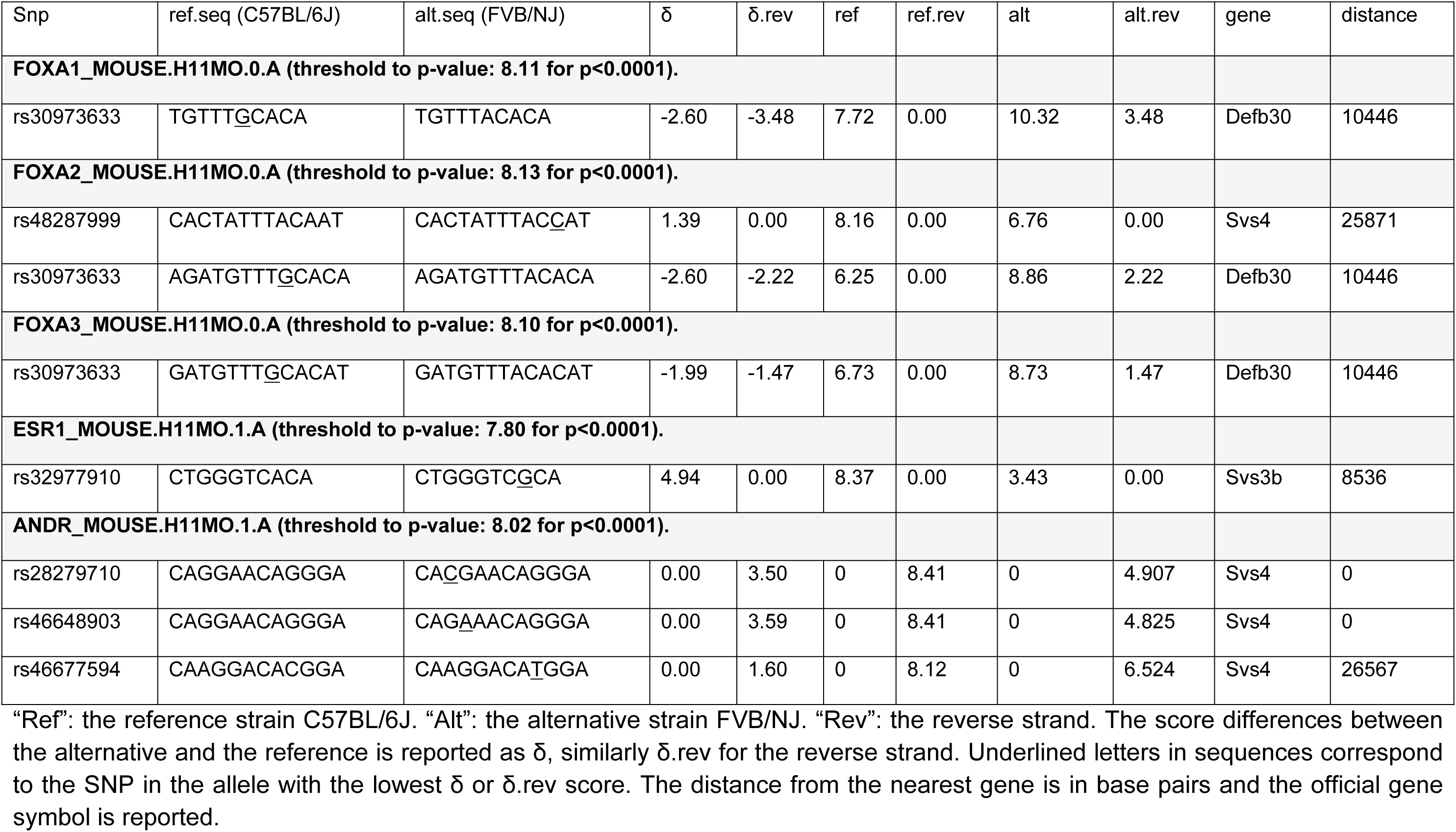
SNP affecting dysregulated targets by disrupting the binding sites for FOXA1-3, ANDR and ESR1.

The second result was the identification of 5 SNP-dependent binding motifs co-localized in the same DNA region encoding the *Svs* genes (Fig 4C). All these SNP-dependent binding motifs were specific to the C57BL/6J allele. More precisely, in the latter, an AR binding site in *Svs4* was affected by two different SNPs (rs28279710 and rs46648903), another AR binding site between *Svs3a* and *Svs6* was affected by rs46677594, an ESR1 binding site upstream of *Svs3b* was affected by rs32977910, and a FOXA2 binding site between *Svs3a* and *Svs6* was affected by rs48287999. All these SNP-dependent binding sites were thus specific of the C57BL/6J allele and all the corresponding genes were silenced by DEHP in C57BL/6J only (Fig 4C and Table 2). This region was defined as the “C57BL/6J susceptibility allele” to prenatal exposure to DEHP.

To summarize, the genome-wide analysis performed resulted in the characterization of the FVB/N resistance allele, involving 4 out of 5 beta-defensins, with all induced by DEHP in FVB/N only, as well as a C57BL/6J susceptibility allele encoding 6 *Svs* genes that were all silenced by DEHP in C57BL/6J only.

### CpG methylation analysis in C57BL/6J-susceptibility allele combined with quantifications of the semenogelins (SEMG) production in sperm

Two independent targeted methods were applied to validate the previous findings on the susceptibility allele of C57BL/6J. First, measures of CpG methylation levels were performed using bisulfite pyrosequencing in the promoters of both *Svs2* and *Svs3ab*. Second, expression levels of SEMG1 and SEMG2 proteins encoded by *Svs2* and *Svs3ab*, respectively, were analyzed by Western blots (Fig 5A). DNA and protein samples required for these analyzes were extracted by sequential precipitations from the TRIZOL interphase and organic phase that remain in frozen sperm samples after RNA extraction.

**Fig 5.**
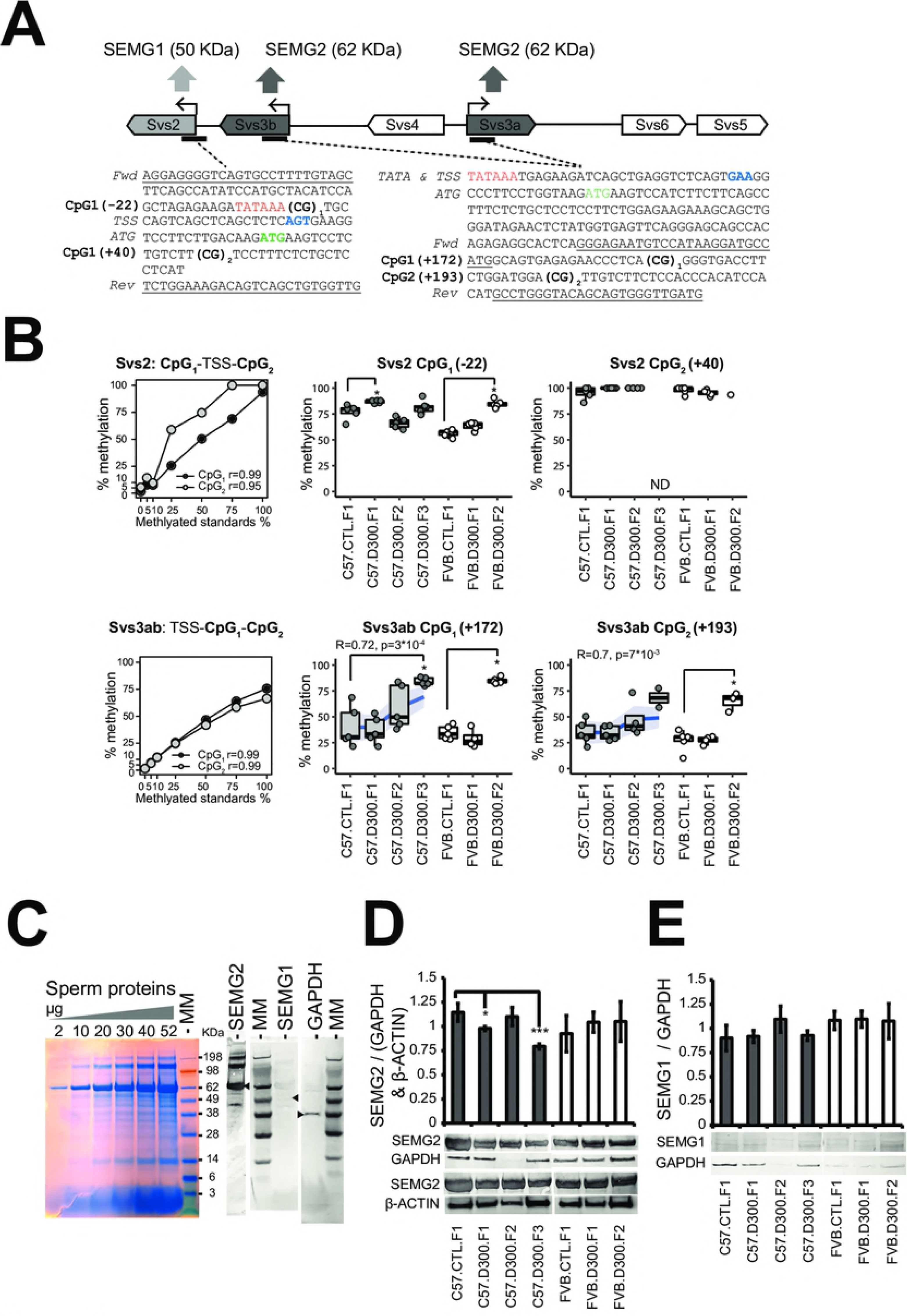
Quantifications of SEMG1-2 combined with measurements of CpG methylation levels in the Svs2 and Svs3ab promoters. (A) Design of bisulfite pyrosequencing assays to measure CpG methylation levels in the promoters of Svs2 encoding SEMG1 and Svs3a/Svs3b encoding SEMG2. Svs3a and Svs3b duplicated-inversed genes are named Svs3ab in part B. The binding sites for the forward and the biotinylated reverse primers used for the bisulfite pyrosequencing experiments are underlined in both original sequences. Two CpG sites were tested in both promoters and numerated CpG1 and CpG2. TSS: transcriptional start site as the +1 coordinate’s reference in blue. ATG: translational starts in green. TATA: tata boxes in red. (B) CpG methylation levels in both Svs2 and Svs3ab promoters were validated using a pre-mixed calibration standard (0, 5, 10, 25, 50, 75 and 100% Fig 5B and then measured in sperm-extracted DNA under each experimental conditions using bisulfite pyrosequencing in quintuplicates. *Increased methylation levels were found in three of four CpG sites of both promoters in both backgrounds in the DEHP exposed filiations compared with their respective controls; Svs2-CpG1 in C57.D300.F1 and in FVB.D300.F2 (both p<10^−2^); Svs3ab-CpG1 in both C57.D300.F3 and in FVB.D300.F2 (both p<10^−3^); Svs3ab-CpG2 in FVB.D300.F2 (p<0.04). Note that methylation levels in the C57BL/6J background correlated positively with the ordered generations in both sites tested in Svs3ab. Each point represents a measure obtained in one mouse. Some measurements did not reach quintuplicates due to full consummation of the DNA samples. (C) Electrophoresis of total sperm protein and Western blots analysis revealed that SEMG2 (62 KDa) is the major protein fraction in sperm. (5) Western blot quantifications revealing decreased SEMG2 in sperm of C57.D300.F1 (p<0.04) and C57.D300.F3 (p<0.004) compared with C57.CTL.F1. Normalization was performed relative to GAPDH (37 KDa) and ϐ-ACTIN (42 KDa). No changes of SEMG2 were recorded in FVB/N. (E) SEMG1 (50 KDa) levels were low and unchanged across the different conditions tested.

The bisulfite pyrosequencing results demonstrated increased methylation levels in three of four CpG sites localized in promoters of both *Svs2 and Svs3ab* in each of the exposed lineages affecting different generations (*Svs2*-CpG_1_ methylation increased in C57.D300.F1 and in FVB.D300.F2; *Svs3ab*-CpG_1_ methylation increased in both C57.D300.F3 and in FVB.D300.F2; *Svs3ab*-CpG_2_ methylation increased in FVB.D300.F2), whereas *Svs2* CpG_2_ was always fully methylated (Fig 5B). Methylation levels in both CpG sites of *Svs3ab* promoter were also increased gradually across the exposed lineage in C57BL/6J as revealed by correlations analyses (Fig 5B). Electrophoresis of total protein extracts combined with Western bolt revealed that SEMG2 is the most expressed proteins in sperm samples, whereas SEMG1 was detected at very low levels (Fig 5C). Western blot quantifications revealed decreased SEMG2 specifically in the exposed linage of C57BL/6J at F1 and F3, without changes in FVB/N (Figure 5D). SEMG1 levels were unchanged across the different conditions tested in both backgrounds (Fig 5E).

Overall, the DEHP-induced increased methylation levels recorded at the promoters of *Svs3ab* in the exposed lineage of the strain C57BL/6J were compatible with the observed decreases of SEMG2 expression in C57.D300.F1 and C57.D300.F3 conditions (Fig 5D). In addition, the results provide a molecular explanation for the DEHP-induced transgenerational decrease in sperm velocities recorded in the C57BL/6J lineages (Fig 1 D, VAP, VCL and VSL). The latter might be due to the transgenerational inheritance of decreased SEMG2 production mediated by increased promoter methylation in *Svs3ab*. Indeed, it is recognized that a fundamental physiological function of semenogelins is the control of sperm motility [107]. DEHP effects on *Svs3ab* in C57BL/6J strain are coherent with the general consensus, showing an increased methylation, a decreased RNA transcription, and a decreased level of the corresponding protein, SEMG2. DEHP effects on *Svs2* in this same strain are also coherent, showing an increased methylation and a decreased RNA transcription. The absence of effect on the corresponding protein, SEMG1, might be due to technical difficulties associated with the low level of this protein. The results obtained in the FVB/N strain were also coherent with the lack of effect of DEHP, except for the increase in both promoter methylation observed in F2 offspring.

## Discussion

C57BL/6J mice prenatally exposed to DEHP present anti-androgenic and pro-estrogenic symptoms, whereas FVB/N mice are phenotypically clearly unaffected. The phenotype in the C57BL/6J strain was clearly affected by prenatal exposure to DEHP on all tested parameters (Fig 1C and 1D). In C57BL/6J, DEHP induced various symptoms in the male reproductive system, reflecting perturbations of sex hormone signaling, such as decreased AGD, a marker of anti-androgenic actions occurring during the prenatal development period [7]. Decreased AGD after exposure to DEHP has been significantly observed in meta-analyses performed in both mice and humans [6]. Decreased sperm counts after prenatal exposure to DEHP are most probably a consequence of alterations in the proliferation of the fetal male germ cell precursors in the gonadal differentiation period at the time of exposure, or may be due to a decrease in the proliferation of Sertoli cells that positively control the germ cell numbers [108]. Similarly, decreased testis weight after prenatal exposure to DEHP may be due to an alteration in the initial proliferation of all or any gonadal cells lineages during the gonadal differentiation period, or may also involve a long-term pro-estrogenic impact. Indeed, the latter, sustained in a long-term manner in the male rete testis increases fluid reabsorption with a consecutive decreased testis weight. In a normal situation, the fluid that surrounds the sperm is subjected to resorption in the efferent ducts by estrogen-dependent actions and in the presence of “enormous amounts” of ER [109]. Interestingly, ER knock-out mice were infertile and showed increased fluid pressure within testes due to the accumulation of the fluid at its production site with decreased sperm counts [110]. Overall, DEHP-mediated anti-androgenic and pro-estrogenic symptoms that remain poorly understood were detected in C57BL/6J, but absent in FVB/N, consistent with an intrinsic resistance of the strain to this recognized endocrine disruptor (Fig 1A and 1B).

In the susceptible C57BL/6J background, initial prenatal exposure to DEHP induced an intergenerational inheritance of anti-androgenic and/or pro-estrogenic symptoms, as well as a transgenerational inheritance of decreased sperm velocity affecting all generations tested up to F3 (Fig 1C and 1D). Importantly, the multigenerational impact found in the C57BL/6J exposed lineage cannot be formally proved in the present study. Indeed, the experimental design lacks F2 and F3 control conditions in both backgrounds, i.e. “FVB.CTL.F2”, “C57.CTL.F2” and “C57.CTL.F3”. This was due to the priority that was given to perform comparisons between both strains.

However, the DEHP-induced transgenerational transmission of decreased sperm velocities recorded in the exposed C57BL/6J seems to be consistent when compared with stable levels recorded for the same parameters in the exposed lineage of the FVB/N strain (Fig 1B compared with Fig 1D). Moreover, the C57BL/6J inbred mouse strain appeared in 1937 and was maintained by mating. How would that be possible if the decreased sperm velocities observed in the C57BL/6J-DEHP-exposed lineage were related to a generational impact independent of the initial DEHP exposure? With some difficulty, we obtained the third generation that showed the strongest decreased sperm velocities after one year. Finally, transgenerational inheritance induced by prenatal exposure to DEHP was previously established for cryptorchidism and testicular germ cell in CD-1 outbred mice and Sprague Dawley rats [111, 112], whereas transgenerational inheritance of altered estrous cyclicity and decreased folliculogenesis were established in female CD-1 mice prenatally exposed to DEHP [113, 114]. These previous reports strongly suggest that DEHP may trigger pro-estrogenic and anti-androgenic effects in a multigenerational manner, at least in some strains of mice and rats.

It has been shown recently that haploid spermatozoa, considered as transcriptionally inert for a long time, carry the RNAs that may influence the development of the embryo by delivering all the paternal RNAs to the egg [115]. Moreover, sperm microRNAs were shown to directly support transgenerational inheritance in mice [116]. For these reasons, we postulate that dysregulated sperm RNAs may be a marker of DEHP exposure and may additionally support the transmission of altered traits to subsequent generations in the sensitive C57BL/6J background. In the present study, the results demonstrated that androgen-regulated, sperm-specific transcripts probably involved in spermatogenesi were affected strain-specifically by prenatal exposure to DEHP (Fig 2). This was not the case for other RNA types, such as microRNA, that were effectively quantified by the all-RNA-seq approach. However, they did not show statistically significant dysregulation across conditions, except for the microRNA mir192, which was included in the 62 dysregulated targets associated with the pattern of generational-dependent changes (Fig 2D and Table 1). Two SNPs affecting SHP differed between both strains and may mediate strain-specific responses to DEHP exposure. Rs387782768 is associated with the absence of FOXA3in FVB/N (Fig 3A) and this explains how DEHP-induced alterations in the expression of FOXA3-dependent genes specifically occur in C57BL/6J, but not in the FVB/N strain. Of note, the strain-specific DEHP effect occurs in the absence of FOXA3 binding site differences between the promoters of C57BL/6J and FVB/N genes (Fig 3C). In addition, the ESR1 gene in the FVB/N strain carries a C to A conversion at rs29315913, affecting the ligand-binding domain of one of the five ESR1 proteins by replacing glycine 447 for a valine [13]. We detected similar levels of ESR1 expression across conditions, but with significantly higher expression levels of the ESR1 transcript variant 4 (NM_001302533) containing the SNP-affected ligand binding domain in FVB/N and an almost absence of its expression in C57BL/6J, independent of DEHP exposure (Fig 3A). Concordantly, a 3-fold higher number of ESR1 binding sites were detected across promoters of genes showing DEHP-mediated induction specific to FVB/N (Fig 3C). Under such circumstances, FVB/N-specific DEHP-induced expression can be explained by the pro-estrogenic impact of DEHP mediated by ESR1 variant 4 in FVB/N in *Gpx5, Spag11b, Lcn8*, *Lcn9*, *Lcn5*, and *Spint4*. Although all contain ESR1 binding sites in their promoters, this does not explain the induction of *Defb30 and Rprl3* in which ESR1 binding sites were not detected (Fig 3C). Of note, the involvement of ER in DEHP-mediated toxicity has been reported previously in DEHP-exposed female mice with reduced ovarian expression of ESR1 in wild type, but not in PPARα(−/−) knockout mice, suggesting that DEHP may induce PPARα mediated repression of ESR1 in the ovary [117].

FVB/N resistance to DEHP is associated with a DEHP-induced expression of various beta-defensins in a resistant allele containing an additional FOXA1-3 binding site due to SNP rs30973633. Transcriptomic results were consistent with a pro-estrogenic property of DEHP in the pool of targets associated with resistance to DEHP in FVB/N (Fig 2C), whose promoters contain higher ESR1 binding sites (Fig 3C). These targets were previously involved in sperm resistance mechanisms (Table 1) against bacterial infections or innate host defense (*Defb30*, *Spag11b*), in protective mechanisms against oxidative stress (*Gpx5*), or in the clearance of exogenous compounds (*Lcn5*, *Lcn9*, *Lcn5*) [37, 118]. Previously, lipocalin 2 (*Lcn2*) was upregulated by 4.6 fold 24 h after exposure to 2 g/kg body weight of DEHP in the testis of male Sprague-Dawley rats [119] and lipocalin 7 was upregulated in C57BL/6J mouse fetal testis following *in utero* exposure to dibutyl phthalate [120]. In addition to these targets, we also identified the serine protease inhibitor Kunitz type 4 (*Spint4*), the cystatin E2 (9230104L09Rik) and the ribonuclease P RNA-like 3 (*Rprl3*) (Table 1). Surprisingly, *Spag11b*, *Spint4*, *Defb30*, *Gpx5, 9230104L09Rik, Lcn5*, *Lcn8* and *Lcn9* were associated with high transcriptional levels despite higher promoter methylation, which seems a paradox when considering that promoter methylation represses gene expression. This role of DNA methylation in gene activation has been reported previously, notably in FOXA2, where the promoter region of the gene displayed also paradoxically high levels of DNA methylation in expressing tissues and low levels in non-expressing tissues [121]. Lack of DNA methylation in the FOXA2 promoter region allowed the binding of the polycomb group repressing proteins, whereas methylation resulted in the loss of repressing proteins in the promoter triggering FOXA2 activation. FOXA2 interacts with the AR to regulate prostate and epididymal genes differentially, particularly lipocalins [122], whereas polycomb directs the timely activation of germline genes in spermatogenesis [123]. Thus, the paradoxical role of promoter methylation may involve repressing proteins.

We also identified a FVB/N-specific binding motif associated with a FVB/N-specific and DEHP-induced induction in loci encoding several β-defensins (Fig 4B). Indeed, various β-defensins have been shown to play a dual role in the sperm, protecting the sperm from bacterial infections, but also acting in a positive manner on sperm motility. Decreased sperm motility was found after CRISPR/Cas9-mediated disruption of various β-defensins (*Defb23*, *Defb26* and *Defb42*), among which only *Defb42* was identified in the resistant allele in our study [124]. Other β-defensins were also found to exert a positive action on sperm motility across various organisms. Human β-defensin 114 protects sperm motility loss due to lipopolysaccharide (LPS) [125]. β-defensin 15 was required for sperm motility in *Rattus norvegicus*, according to experiments involving co-cultures of spermatozoa with *Defb15* antibody, lentivirus-mediated RNA interference (RNAi) and knock down of *Defb15* [126]. Bovine β-defensin 126 promoted sperm motility in cattle according to experiments involving sperm treated with a recombinant protein [127]. The β-defensin “SPAG11E” protected sperm motility loss after LPS treatment in the rat [128]. In humans, the β-defensin 1 was also implicated in sperm motility in studies where interference with the *Defb1* function decreased sperm motility, which was able to be restored by recombinant *Defb1* [129]. In the rat, the β-defensin Bin1b, homologous to human β-defensin 1, induces progressive sperm motility in immotile immature sperm [130].

Multiple SNPs (rs28279710, rs46648903, rs46677594, rs32977910, rs48287999) affect sexual hormone signaling in the C57BL/6J-susceptible allele encoding the *Svs* genes. Targets for DEHP-susceptibility were found in C57BL/6J mice (Fig 2E, red lines; Fig 3B and 3C, red points/symbols) and previously associated with a TDS phenotype [20]. They include the androgen-dependent *Saa2*, the serine peptidase inhibitor Kazal type 1 *Spink1*, the poorly described *9530003J23Rik*, as well as the androgen-dependent *Sva*, *Svs2*, *Svs3a*, *Svs3b*, *Svs4*, *Svs5*, *Svs6* and *Pate4* genes (Table 1). Previous independent reports showed a downregulation of S*vs5* by a minus 3.75-fold change in microarray. This was confirmed by RT-qPCR in the fetal testes of Sprague-Dawley outbred CD rats exposed *in utero* to 750 mg/kg per day of DEHP during E12-19 in association with reduced AGD, and also when exposure consisted of dibutyl phthalate and two other phthalates [131]. Targets were silenced by methylation of their promoters (Fig 3B) that contained AR binding sites, notably in *Sva*, *Svs2* and *Svs3ab* (Fig 3C). In addition, the targets exerted a direct impact on sperm physiological functions, notably by regulating sperm motility (Table 1). In our study, the analysis revealed SNPs affecting sexual hormone-binding sites specific to C57BL/6J and co-localized in the DNA encoding *Svs2*, *Svs3ab*, *Svs4*, *Svs6* and *Svs5* genes. The SNPs may explain the strain-specific DEHP response by rendering the allele more responsive to SHP in C57BL/6J compared with FVB/N (Fig 4C). For this reason, DNA regions encoding the *Svs* genes in C57BL/6J and affected by SNP represent a susceptible allele.

The C57BL/6J-specific decrease in sperm velocity induced by *in utero* exposure to DEHP is associated with both silencing of the *Svs3ab* gene and the decreased production of SEMG2. Among genes encoded by the susceptible allele, both *Svs2* and *Svs3ab* encode murine Semenogelins SEMG1-2, which represent the most abundant protein fraction in sperm, as well as regulating sperm motility [132]. In parallel, we discovered an apparent DEHP-induced transgenerational inheritance of decreased sperm velocity measured with three different parameters (VCL, VAP and VSL), specifically affecting the C57BL/6J background and tested up to the third filial generation in its exposed lineage (Fig 1D). Western blot and bisulfite pyrosequencing results revealed that the transgenerational sperm motility decrease was probably mediated by increased methylation in CpG sites of *Svs3ab* promoters by affecting SEMG2 production in a transgenerational manner, specifically in the C57BL/6J strain (Fig 5).

In conclusion, prenatal exposure to the anti-androgenic DEHP molecule induces intergenerational and transgenerational alterations observed in a susceptible mice background, but not in a resistant mice strain. We found converging evidence for strain-specific, SNP-mediated alterations in androgen-regulated sperm transcripts in association with the alteration of several male fertility parameters transmitted across several generations in C57BL/6J. This transmission is mediated by the epigenetics of the paternal DNA having escaped the erasure-reprogramming system or by other spermatozoa-dependent unknown mechanisms. Biological support for transmission across generations may also involve dysregulated sperm RNA. However, injections of sperm RNA into fertilized oocytes, technically difficult, would be required to formally prove their transmission of the phenotype. We observed that strain-specific genomic variations may confer complex resistance mechanisms. SNPs were identified as associated with the strongest DEHP-induced specific changes in the germ cell content of RNAs coding for proteins required for sperm motility. First, rs30973633 was shown to add a FOXA1-3 putative binding motif in the FVB/N allele encoding various β-defensins overexpressed by DEHP specifically in that strain, with a positive impact of DEHP exposure on FOXA1-3 transcript levels. Various SNPs (rs32977910, rs28279710, rs46648903, rs48287999 and rs46677594) that decrease the androgenic regulation of the *Svs* genes in the FVB/NJ allele may be associated with DEHP-induced epigenetic silencing in C57BL/6J. Finally, the relevance of these findings for humans remains to be investigated. By extrapolation, the results suggest that human embryos exposed to DEHP may not be equally affected in their future reproductive health. This study shows how mouse genomic variations may be associated with complex resistance mechanisms to prevent the inter- and transgenerational inheritance of altered male fertility parameters. More fundamentally, this study presents a good model to better characterize the array of endocrine signaling and target genes involved in the male reproductive function.

## Materials and methods

This study was approved by the Ethics Committee for Animal Experimentation of the University of Geneva Medical School (Geneva, Switzerland) and by the Geneva Cantonal Veterinarian Office (permit reference: G61/3918) under the licenses GE/9/15 from February 2015 to August 2016 and GE/115/16 since 2017.

All resources used in the present work are given with the Resource Research Identifiers (RRID) number when possible, as recommended by NIH guidelines (https://scicrunch.org/). All the metadata generated in the study were deposited in a publically-available registry for transparency purposes. The data have also been deposited in the NCBI Gene Expression Omnibus [133] as GEO series accession number GSE107839. Files were converted by NCBI to Sequence Read Archive (SRA) files. We reused 10 samples from the previous GEO series accession number GSE86837. The MBD-seq data were reused from the GEO series accession number GSE67159. The analysis of SNPs impacting on hormonal binding motifs in the function of strain-specific RNA responses in male germ cells was deposited in the Mendeley repository (https://data.mendeley.com/datasets/3s94xbbtjx/draft?a=10be535a-1801-4805-977a-1b8d83b058f7).

### Experimental mice model

C57BL/6J (Charles River France, RRID:IMSR_JAX:000664) and FVB/N (Charles River Germany, RRID:IMSR_CRL:207) mice were maintained at the animal Core Facility of the University of Geneva Medical School in standard plastic housing cages with *ad libitum* access to food (RM3, SDS Dietex, France) and water, with a 12:12 light cycle. Table S3 presents the ARRIVE checklist for this study.

### Prenatal exposure to DEHP and filial generations

One male and one female were mated in the same cage during the night and the presence of a copulatory plug the next day was considered as embryonic day 1 (E1). Prenatal exposure was performed daily from E9-19 covering the sexual differentiation period of the embryos by *per os* injections of 20 μl fixed volume to restrained pregnant female mice. Corn oil delivery vehicle for fat-soluble compounds (Sigma, C8267) was injected in controls (CTL) and 1.15 M DEHP (Fluka-Sigma, DEHP, 80030) diluted in corn oil was injected in exposed mice (D300). The dose of DEHP (CAS No. 117-81-7) was calculated for an estimated mouse weight of 30 g and corresponded to 300 mg of DEHP per kg of mice per day. F1 males prenatally exposed to DEHP were crossed with newly-ordered females of the respective backgrounds to produce F2. The third filial generation (F3) was obtained in the C57BL/6J background only by crossing F2 males with newlyordered females. The experimental design is shown in Fig 1A and 1C.

### Computer-assisted sperm analysis and RNA extraction from spermatozoa

Sperm samples were extracted from 100 days of life age-standardized males. The left and right cauda epididymides of each mouse were cleanly dissected and deposited in a petri dish (Falcon^®^, 351008) filled with a pre-warmed 1 ml M2 medium (Sigma, M7167). Epididymides were incised to allow the sperm to swim out during 10 min at 37°C. Sperm suspensions were transferred in a 100 µm-deep homemade chamber placed on a stage warmer to maintain the temperature at 37°C during the CASA process (Minitherm^®^, Hamilton Thorme Lac, Beverly, MA, USA). Observations were made with a 4X phase contrast Objective (Olympus, RMS4X) at a final magnification of 40x with fixed parameters. For each sample, a minimum of 400 sperm cells tracks were captured at a rate of 25 images per sec. In parallel, 900 µl of sperm suspension were collected without tissue debris and centrifuge to pellet the sperm. RNA extraction was then performed using TRIZOL according to the manufacturer’s recommendations and including glycogen as carrier (Invitrogen, UltraPure™ Glycogen, Carlsbad, CA, USA). Elution was in 20 µl water (Bioconcept, Water for Molecular Biology DNA/DNAse/RNAse free) and samples were snap-frozen and conserved at −80°C.

### Estimation of the purity of spermatozoa samples

The purity of spermatozoa used to extract RNA was assessed before the procedure by visualizing sperm under a contrast-phase microscope during the CASA process. After RNA sequencing, the purity of sperm RNAs was assessed by read length distributions as corresponding to the profile of mature sperm RNA signatures, involving a peak at 22 for microRNA and a peak at 32 nucleotide for tRNA-derived small RNAs [134]. The read length distribution of all samples is reported in Fig S1.

### Preparation of all-RNA-seq libraries and sequencing

All-RNA-seq was performed as previously described [20]. RNA molecules were treated to be compatible with adapter ligation following a protocol developed by Fasteris SA (Geneva, Switzerland), resulting in RNA molecules carrying a 5’-monophosphate extremity and a free 3’-hydroxyl group independent of their starting differences (Fig 2A). Libraries were prepared using the TruSeq small RNA kit (Illumina Inc., San Diego, CA, USA) and sequenced on a HiSeq 2500 (Illumina Inc.). Base calling was performed using HiSeq Control Software 2.2.58, RTA 1.18.64.0 and CASAVA-1.8.2.

### All-RNA-seq reads mapped to mm10 genome and differential quantification

Fastq files were processed on a Linux workstation (Ubuntu 14.04 LTS) and on the “Baobab” high performance computing cluster of the University of Geneva. The analysis was performed using TopHat and Cuffdiff [135]. Quality control of the sequencing process was performed with fastqc ensuring >Q28 in the quality control report for all reads. RA3 adapter sequence was clipped from fastq files and the minimum reads size was settled at 18 pb. Reads were mapped to the mm10 mouse genome using TopHat with appropriate strand-specificity options activated ($ tophat - p 16 -G genes.gtf --library-type fr-secondstrand -o genome reads.fastq). BAM files were indexed and sorted with samtools and reads statistics were computed to summarize the mapping process (Table S1). Quantifications were performed using TopHat.

### Hierarchical cluster analysis and heat map representation

Hierarchical cluster analysis was performed in R with iterative tests involving the diffData function of CummeRbund [136]. A two-dimensional graphical false-color image representation (heatmap) of the gene expression data was produced in R using the heatmap.2 function of the gplots package (Fig 2B) [137].

### Analysis of pathways

Selected genes were tested for enrichments using STRING [104, 138]. The analysis was performed online (http://string-db.org/) with the list of official gene symbols and selecting the “*Mus musculus*” organism. The minimum required interaction score was set at the highest confidence level (0.9). A significant enrichment was defined based on a false discovery rate (FDR) <0.01. The representative enrichment per category are reported in Table S2.

### Expression of identified targets in germ cell subtypes

Expression of genes across male germ cell subtypes came from a previously published independent study [12]. Primary spermatocytes and haploid round spermatids were isolated from B6129SF2/J mice at 8-9 weeks, whereas spermatogonia were isolated from testis of 4-8 days postpartum mice using enzymatic digestions of decapsulated testis combined with sedimentation techniques. In our previous study, enriched germ cell populations were analyzed using the microarrays “A-AFFY-45” (Affymetrix GeneChip Mouse Genome 430 2.0). The data were deposited in the ArrayExpress database archived in the European Bioinformatics Institute under the accession number “E-TABM-130”. Data were uploaded and integrated in our analysis, notably for Fig 2G.

### Combined promoter methylation and RNA expression analysis

All-RNA-seq and MBD-seq data available for all four lineages at F1 were merged by gene symbol in R software separately for both strains (Fig 3B). For each gene, the promoter is defined as the region spanning 2000 bp upstream of the gene’s 5’-end to 200 bp downstream. Its methylation level is measured as the number of reads obtained in a 2.2 kilobase probed region divided by the number of CpG in the tested region [10].

### Detection of binding sites for the sexual hormone signaling performed in the promoters of the identified targets

Binding sites for FOXA1-3, ESR1 and AR were detected in promoters from −2000 to 0 bp among the selected targets using Rsat [105] (Fig 3B). Briefly, promoter sequences were extracted from *Mus musculus* GRCm38 using a list of official gene names and an analysis was performed using the matrix-scan full option with the same HOCOMOCO position weight matrices (PWM) used afterwards [139].

### Genomic variation between FVB/N and C57BL/6J strains

Sequencing and characterization of the FVB/N strain compared with the reference C57BL/6J strain was performed by others [13]. This study generated a catalogue of the major genomic variations specific to FVB/NJ strains corresponding to the FVB/N strain imported into the Jackson Laboratory. When possible, SNPs were verified between both strains analyzed (FVB/N and C57BL/6J) using the all-RNA-seq mapped reads on the IGV and showed concordant genotypes compared with the FVB/NJ and C57BL/6J strains in Wong et al [13] (Fig S2).

### SNP-dependent hormonal binding motifs analyzed in function of strain-specific RNA responses to DEHP exposure

Bioinformatic approaches were used to assess strain-specific SNP affecting binding sites for hormonal controls and genes strain-specifically dysregulated by prenatal exposure to DEHP on genome-wide scale. The R packages and libraries “VariantAnnotation” [140], “Biostrings” [141], “BSgenome.Mmusculus.UCSC.mm10” [142] and “EnsDb.Mmusculus.v79” [143] were required for the analysis. 5’556’605 FVB/NJ SNP variations compared with C57BL/6J were uploaded from the FTP site of the Sanger Institute and filtered to 4’823’906 of high quality. Filtrated SNPs between both strains were used to reconstruct C57BL/6J and FVB/NJ strain-specific alleles with sizes of sequences equal to two times the size of HOCOMOCO position weight matrices (PWM) for mouse AR (named ANDR in the HOCOMOCO PWM and replaced by AR in the present text for simplification purpose), ESR1 and FOXA1-3 [139]. The “matchPWM” function in R was used to computed strain-specific identification scores based on the SNP and the respective scores were reported for the reference (C57BL/6J = ref) and the alternative (FVB/NJ = alt) alleles in both the reverse complement and the positive strand [144]. The results were annotated reporting the distance and the name of the most proximal genes from the SNP-dependent motifs. Genome-wide statistical significances of SNP-dependent detected motifs were based on the HOCOMOCO threshold to p-value relationships. Tables were created with the resulting detected SNP-dependent motifs. Generated tables were then merged individually to the all-RNA-seq derived FPKM data and deposited in Mendeley. SNPs affecting the expression of genes in a strain-specific manner by abolishing or creating a binding motif for FOXA1-3, AR and ESR1 were identified using δ, for motif specificity, in function of Z, reflecting strain-specific transcriptional changes; (δ <−1 or > 1, [δ = score (C57BL/6J) – score (FVB/NJ)], Z< −7 or > 7, 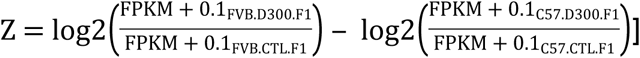. The genome-wide significant results are reported in Table 2.

### Validation experiments involving bisulfite-pyrosequencing

DNA was extracted using ethanol precipitation from the organic phase of TRIZOL conserved at −20°C. These samples correspond to the sperm samples obtained during the CASA process that were used for RNA extraction. Bisulfite conversion of DNA was performed using the EZ DNA methylation-Lightning™ kit (Zymo Research, Irvine, CA, USA; D5030). All assays were validated using methylation standards (EpigenDX, Hopkinton. MA, USA; 80-8060M-PreMix). PCR cycling conditions were 95°C for 15 min, then 50 cycles of 94°C for 30 s, 54°C for 30 s (replaced by 56°C for 30 s for *Svs3b*), 72°C for 30 s, with a final elongation during 1 min at 72°C. Pyrosequencing was performed using a Pyromark Q24 instrument and a workstation from Qiagen (Hilden, Germany) using appropriated enzyme, substrate, nucleotides and reagents. For methylation measured in the *Svs2* promoter, a 175 bp amplicon encompassing 2 CpG sites was generated using forward 5’-AGGAGGGGTTAGTGTTTTTTGTAGT-3’ and biotinylated reverse 5’-[BIO] CAACCACAACTAACTATCTTTCCAAA-3’ primers. The sequencing primer was 5’-GTTATATTTAGTTAGAGAAGATATAAA. The sequence to analyze was YGTGTTAGTTAGTTTAGTTTTTAGTGAAGGTTTTTTTTGATAAGATGAAGTTTTTTGTTTTYGTTTTTTTTTTGTTTTTTATTTTGGAAAGATAGTTAGTTGTGGTTG and the dispensation order was GTCGTGTAGTAGTAGTTAGTGAGTTGACTAGATGAGTTAGTTCTGTTG. The two CpG sites were validated by methylation standards. For methylation measured in the *Svs3b* gene, a 118 bp amplicon encompassing 2 CpG sites was generated using the forward primer Svs3b_F 5’-GGGAGAATGTTTATAAGGATGTTATG-3’ and the reverse Svs3b_R_biot 5’-[BIO]CATCAACCCACTACTATACCCAAAC3’. The sequencing primer was the forward primer. The sequence to analyze was GTAGTGAGAGAATTTTTTAYGGGGTGATTTTTTGGATGGAYGTTGTTTTTTTTATTTATATTTATATGTTTGGGTATAGTAGTGGGTTGATG and the dispensation order was AGCTAGTGAGAGATTGATCGGTGATTGATGTATCGTG. Both CpG sites were validated using methylation standards.

### Western blots of SEMG1 and SEMG2

Sperm total protein samples were obtained from TRIZOL fractions after RNA and DNA extractions according to the manufacturer’s recommendations by precipitation with 100% isopropanol. Protein pellets were washed with 0.3 M guanidine hydrochloride in 95% ethanol and re-suspended in 200 µl of 0.5% SDS 4 M urea by overnight rotations. Total protein was quantified using the Pierce protein assay 660nm (ThermoFisher Scientific, Waltham, MA, USA; réf. 22662) on a nanodrop ND-100 and using the ionic detergent compatibility reagent (IDCR) (ThermoFisher; réf. 22663). Approximately 20 µg of protein per conditions (C57.CTL.F1, C57.D300.F1, C57.D300.F2, C57.D300.F3, FVB.CTL.F1, FVB.D300.F1 and FVB.D300.F2) were separated on precast Bolt TM 4-12% Bis-Tris gel (ThermoFisher; NW04120) at 150 volts during 45 min and transferred to a nitrocellulose membrane (ThermoFisher, iBlot^®^ 2NC Regular Stacks, IB23001) using program P3 on the iBlot2 transfer device (ThermoFisher; IB21001). Membranes were blocked in the SuperBlock^®^ (PBS) Blocking Buffer (ThermoFisher; 37515) during 1 h. Five washes of 5 min each were performed between the incubations with Tween-containing TRIS buffer (TBST, ThermoFisher; 28360). Membranes were then incubated 1 h with antibodies diluted in the SuperBlock^®^ (PBS) Blocking Buffer. Antibody dilutions were 1:5000 α-SEMG2 polyclonal rabbit anti-mouse [antibodies-online.com, ABIN252161], 1:500 α-SEMG1 polyclonal rabbit anti-mouse [antibodies-online.com, ref ABIN2406262], 1:10,000 α-GAPDH monoclonal rabbit anti-mouse-rat-chicken-human [Abcam, ab181602], before incubation with 1:10,000 secondary antibody goat anti-rabbit IgG alkaline phosphatase secondary (antibodies-online.com, ABIN616373). Revelations were performed with NBT/BCIP Ready-to-Use Tablets (Sigma, c.o. Merck, Darmstadt, Germany, catalogue number: 11697471001). Additionally, membranes were incubated with beta-ACTIN monoclonal antibody coupled to HRP (Sigma, A3854) followed by detection using the ECL™ Prime Western blot reagents kit (Amersham, Little Chalfont, UK; RPN2232) with images acquired in the myECL™ Imager with the chemiluminescence mode (ThermoFisher), as an alternative to detect issue with abnormally low levels of GAPDH in condition C57.D300.F2. All images were analyzed using the open source imageJ software.

### Statistical analysis

One-way analysis of variance with post-hoc Tukey honestly significant difference test (ANOVA-Tukey HSD) was used to identify means that were significantly different between conditions for the parameters measured with CASA, as well as for AGD and testes weight measurements (Fig 1). In all-RNA-seq analysis, the CuffDiff computed false-discovery rate adjusted p-values (q-values) were used to identify the transcript levels statistically significantly different between the labeled conditions. Probability estimates for each cluster resulting from the hierarchical cluster analysis were obtained using Pvclust [145]. Pearson’s chi-square goodness-of-fit test was computed in R version 3.4.2 using the “chisq.test” function to assess the statistical significance of differences in frequencies of the respective proportions of genes whose expression increases during spermatogenesis among the identified targets versus all the genes. Correlations between MBD-seq and RNA-seq data were analyzed using non-parametric Kendall’s tau and Spearman’s rho rank correlations in R (Fig 3B). Non-parametric pairwise Wilcoxon test with Benjamini & Hochberg correction for multiple testing was used to estimate differences of transcript levels between conditions associated with SNPs in Figs 3A and Fig 4 [146]. Parametric Pearson correlations between CpG methylation and the ordered conditions involving C57BL/6J strain were performed in R (Fig 5B).

## Acknowledgments

This research was supported by the Swiss Centre for Applied Human Toxicology and the Ernst and Lucie Schmidheiny Foundation. Computations were performed at the University of Geneva on the Baobab cluster. We are grateful to Yann Sagon for his helpful recommendation to use the Baobab cluster. We are also grateful to Jessica Escoffier and Beatrice Conne for their help. We thank Fasteris SA (Geneva, Switzerland) and NXT-DX (Ghent, Belgium) for the preparation and sequencing of the all RNA-seq and MBD-seq libraries.

## Declaration of competing financial interests

The authors declare no actual or potential competing financial interests.

## Supporting information

**Fig S1.** Read lengths distribution across samples involving a peak at 22 nucleotides, specific to the size of mature microRNAs, a peak at 32 nucleotides, specific to the size of tRNA-derived small RNAs, and a peak at 50 bp, involving mainly coding RNAs.

**Fig S2.** Example of SNP validation with all RNA-seq reads in a region of interest: rs28279704 missense mutation validated in the *Svs*4 gene as being “A” in C57BL/6J, reversely in the ATG codon for methionine, and “G” in FVB/N, reversely in the CG codon for threonine.

**Table S1:**
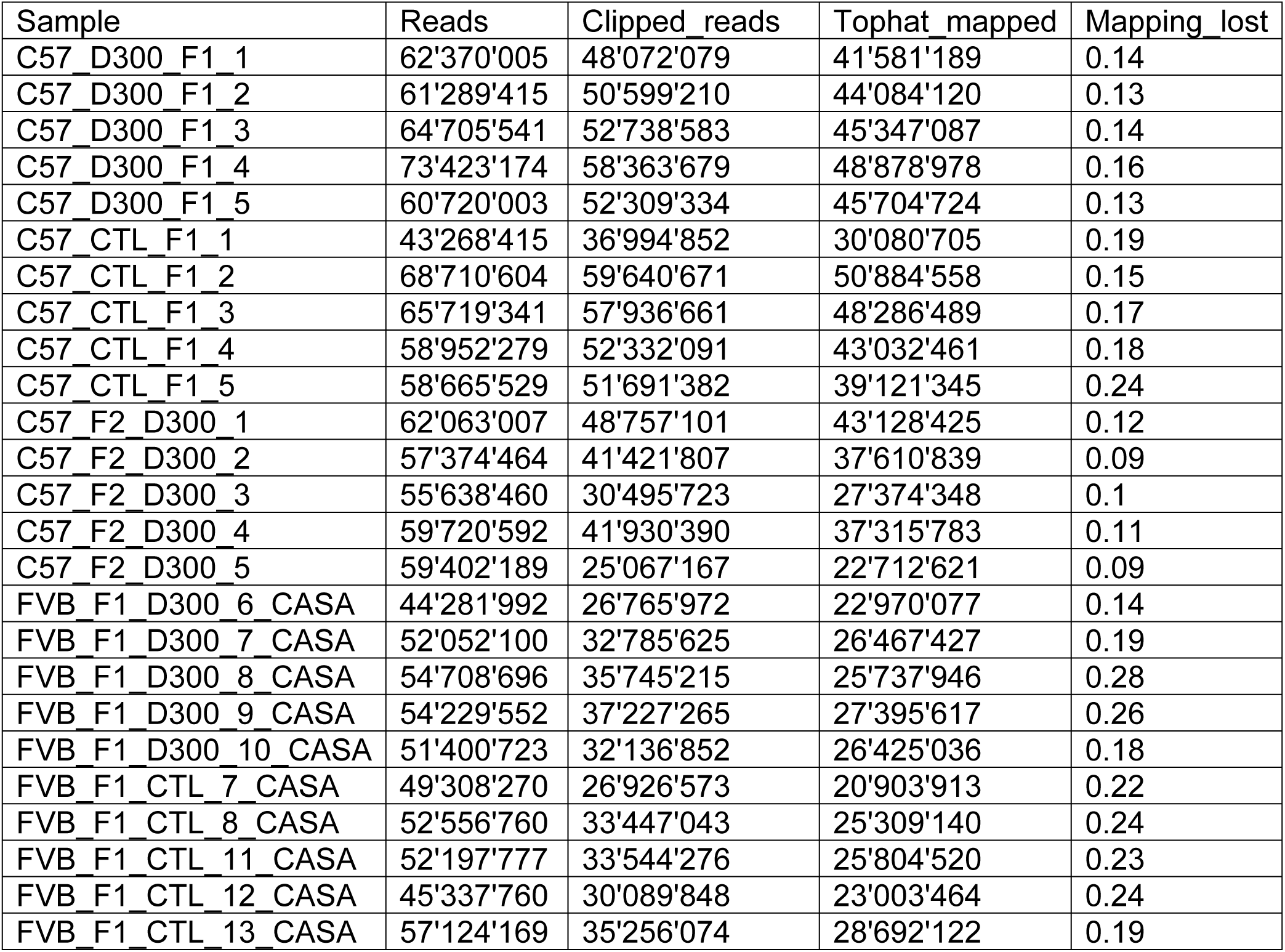
Read log sheet of all RNA-seq.

**Table S2.**
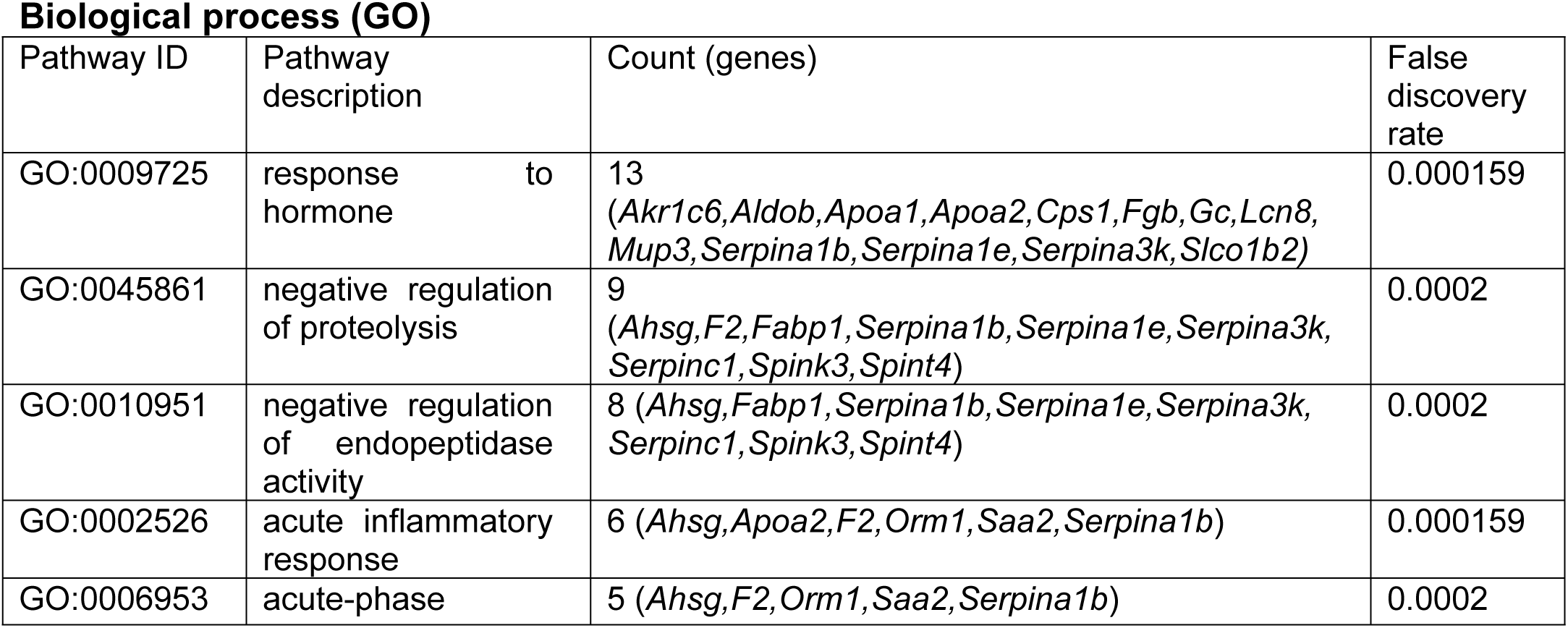
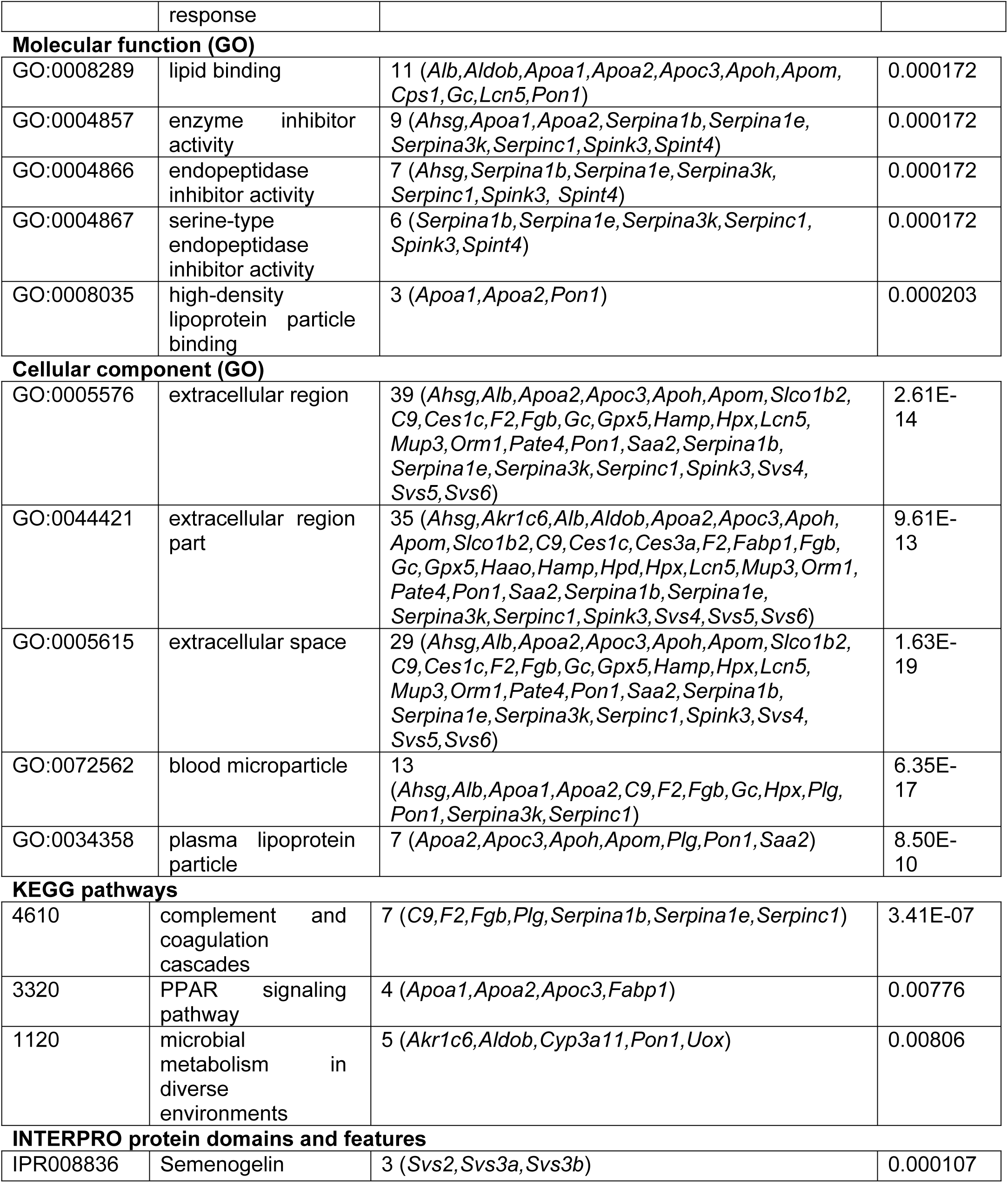
Pathway analysis of the 62 dysregulated RNAs. The statistically significant enrichments in GO terms, KEGG pathways and INTERPRO protein domains identified in the 62 dysregulated sperm RNAs according to the STRING database [104].

**Table S3.**
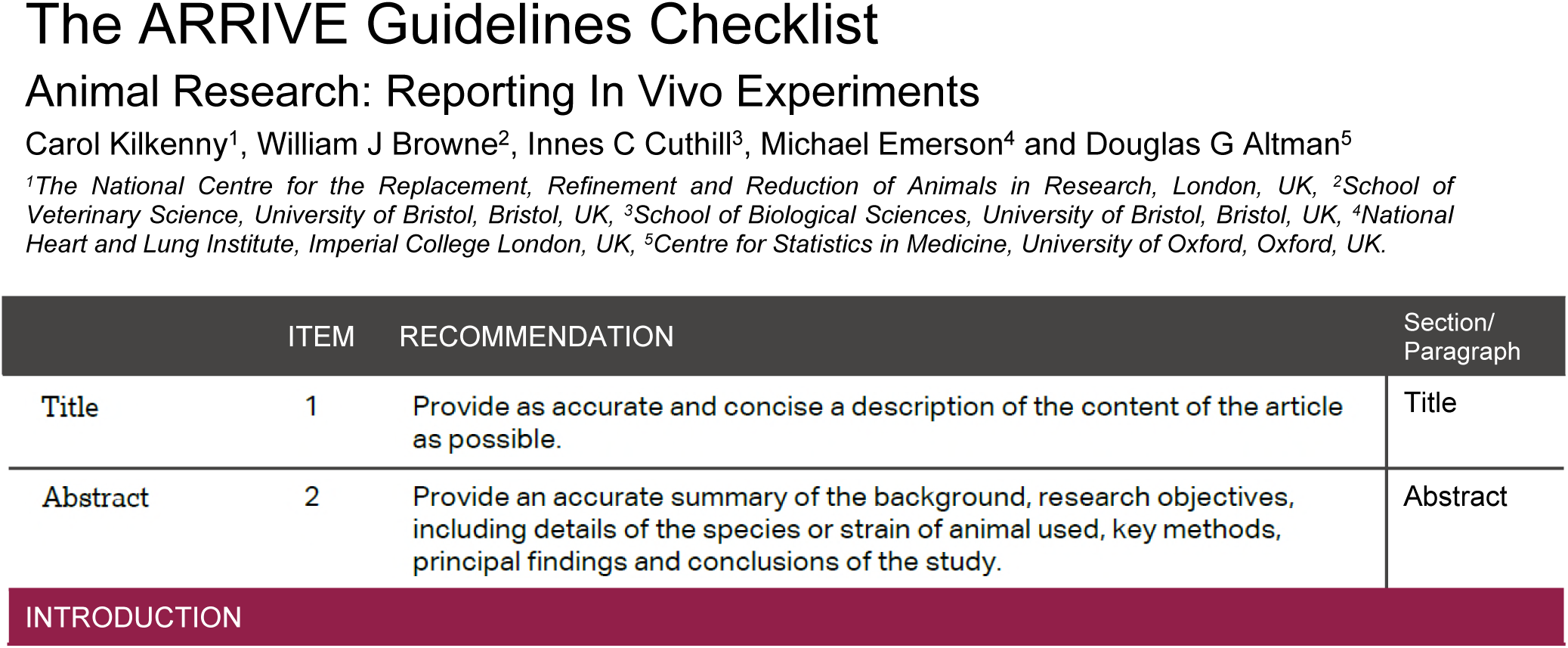
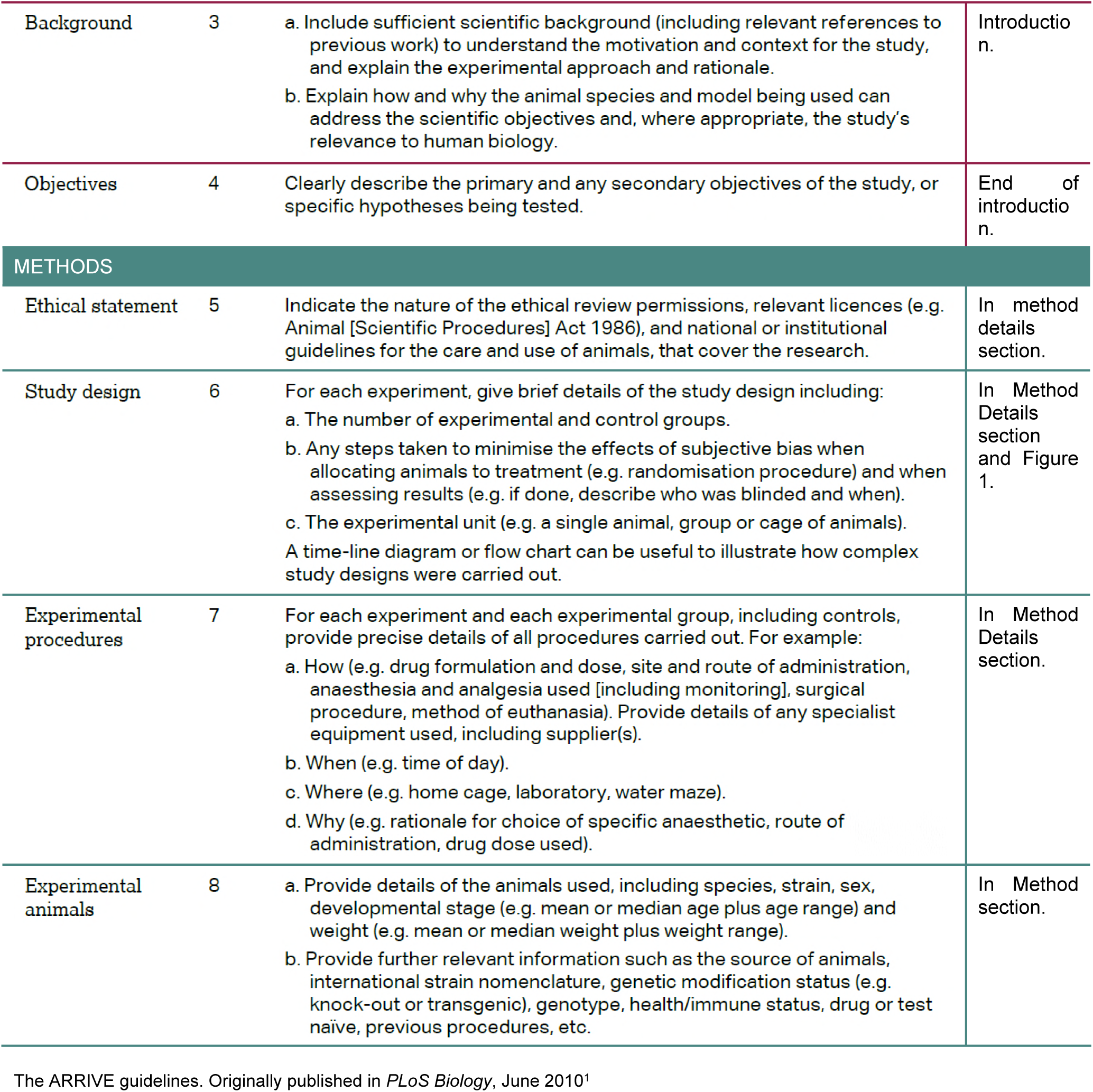
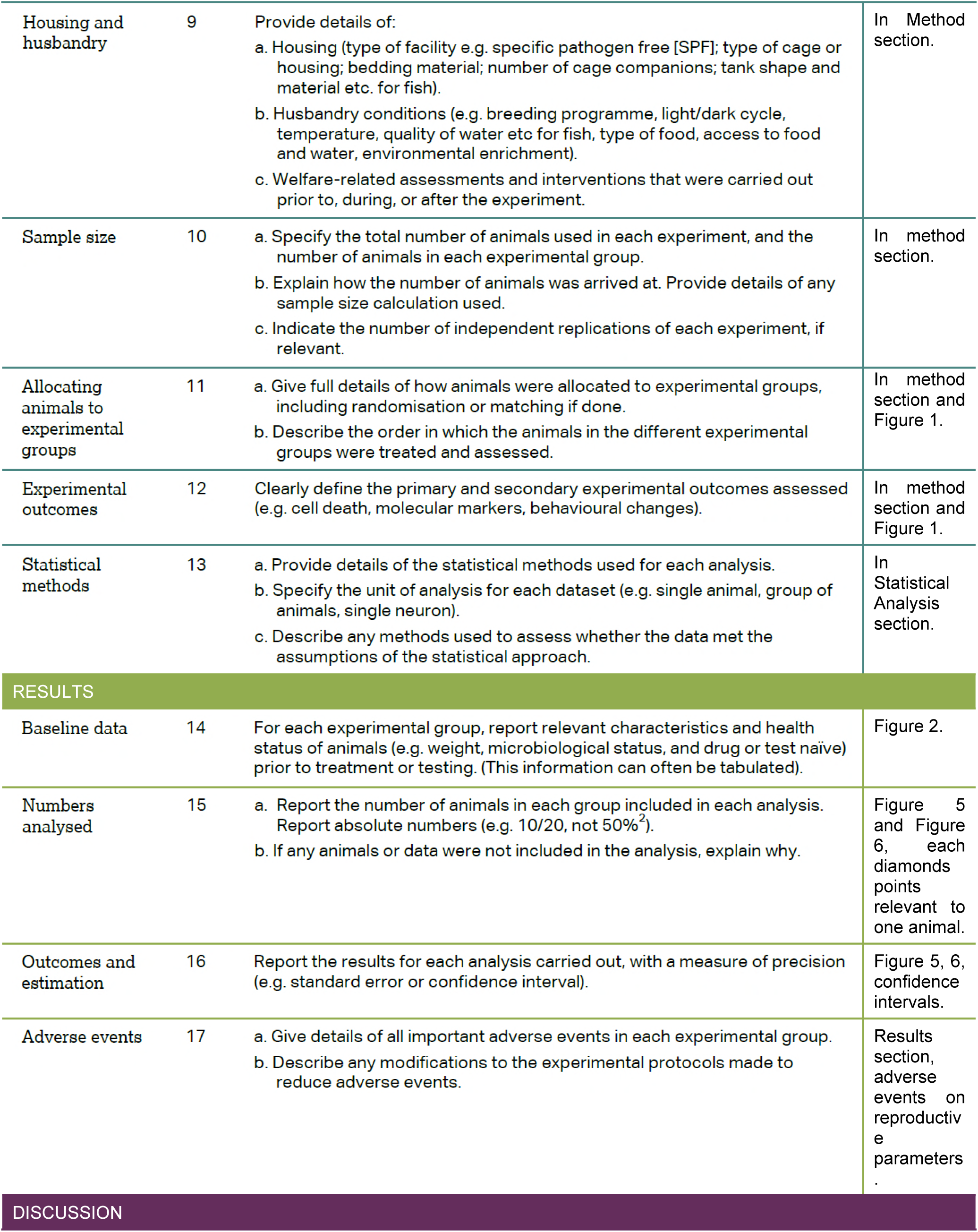
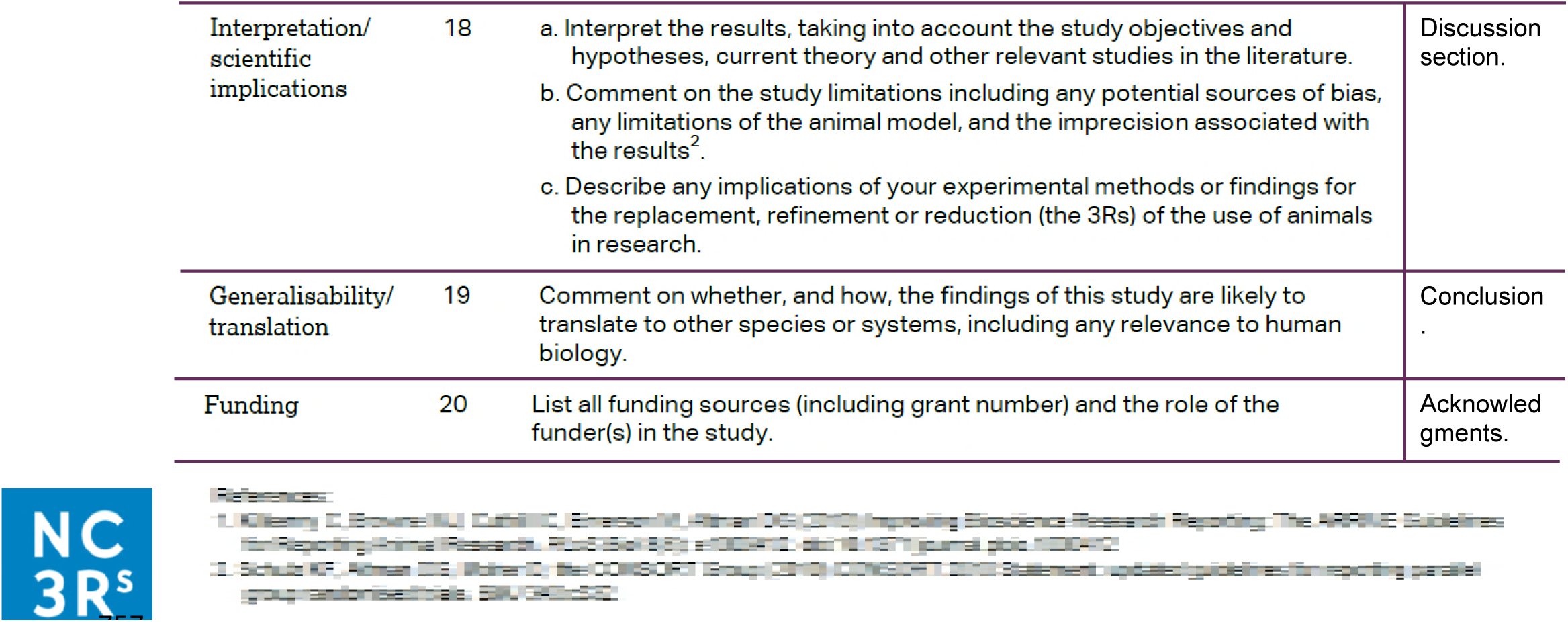
The ARRIVE Guidelines Checklist for reporting animal data completed for this study.

|  | ITEM | RECOMMENDATION | Section/<br>Paragraph |
| --- | --- | --- | --- |
| Title | 1 | Provide as accurate and concise a description of the content of the article as possible. | Title |
| Abstract | 2 | Provide an accurate summary of the background, research objectives, including details of the species or strain of animal used, key methods, principal findings and conclusions of the study. | Abstract |
| INTRODUCTION |  |  |  |
| Background | 3 | a. Include sufficient scientific background (including relevant references to previous work) to understand the motivation and context for the study, and explain the experimental approach and rationale.<br>b. Explain how and why the animal species and model being used can address the scientific objectives and, where appropriate, the study's relevance to human biology. | Introduction. |
| Objectives | 4 | Clearly describe the primary and any secondary objectives of the study, or specific hypotheses being tested. | End of introduction. |
| **METHODS** |  |  |  |
| Ethical statement | 5 | Indicate the nature of the ethical review permissions, relevant licences (e.g. Animal [Scientific Procedures] Act 1986), and national or institutional guidelines for the care and use of animals, that cover the research. | In method details section. |
| Study design | 6 | For each experiment, give brief details of the study design including:<br>a. The number of experimental and control groups.<br>b. Any steps taken to minimise the effects of subjective bias when allocating animals to treatment (e.g. randomisation procedure) and when assessing results (e.g. if done, describe who was blinded and when).<br>c. The experimental unit (e.g. a single animal, group or cage of animals).<br>A time-line diagram or flow chart can be useful to illustrate how complex study designs were carried out. | In Method Details section and Figure 1. |
| Experimental procedures | 7 | For each experiment and each experimental group, including controls, provide precise details of all procedures carried out. For example:<br>a. How (e.g. drug formulation and dose, site and route of administration, anaesthesia and analgesia used [including monitoring], surgical procedure, method of euthanasia). Provide details of any specialist equipment used, including supplier(s).<br>b. When (e.g. time of day).<br>c. Where (e.g. home cage, laboratory, water maze).<br>d. Why (e.g. rationale for choice of specific anaesthetic, route of administration, drug dose used). | In Method Details section. |
| Experimental animals | 8 | a. Provide details of the animals used, including species, strain, sex, developmental stage (e.g. mean or median age plus age range) and weight (e.g. mean or median weight plus weight range).<br>b. Provide further relevant information such as the source of animals, international strain nomenclature, genetic modification status (e.g. knock-out or transgenic), genotype, health/immune status, drug or test naïve, previous procedures, etc. | In Method section. |
| The ARRIVE guidelines. Originally published in *PLoS Biology*, June 2010<sup>1</sup> |  |  |  |
| Housing and husbandry | 9 | Provide details of:<br>a. Housing (type of facility e.g. specific pathogen free [SPF]; type of cage or housing; bedding material; number of cage companions; tank shape and material etc. for fish).<br>b. Husbandry conditions (e.g. breeding programme, light/dark cycle, temperature, quality of water etc for fish, type of food, access to food and water, environmental enrichment).<br>c. Welfare-related assessments and interventions that were carried out prior to, during, or after the experiment. | In Method section. |
| Sample size | 10 | a. Specify the total number of animals used in each experiment, and the number of animals in each experimental group.<br>b. Explain how the number of animals was arrived at. Provide details of any sample size calculation used.<br>c. Indicate the number of independent replications of each experiment, if relevant. | In method section. |
| Allocating animals to experimental groups | 11 | a. Give full details of how animals were allocated to experimental groups, including randomisation or matching if done.<br>b. Describe the order in which the animals in the different experimental groups were treated and assessed. | In method section and Figure 1. |
| Experimental outcomes | 12 | Clearly define the primary and secondary experimental outcomes assessed (e.g. cell death, molecular markers, behavioural changes). | In method section and Figure 1. |
| Statistical methods | 13 | a. Provide details of the statistical methods used for each analysis.<br>b. Specify the unit of analysis for each dataset (e.g. single animal, group of animals, single neuron).<br>c. Describe any methods used to assess whether the data met the assumptions of the statistical approach. | In Statistical Analysis section. |
| **RESULTS** |  |  |  |
| Baseline data | 14 | For each experimental group, report relevant characteristics and health status of animals (e.g. weight, microbiological status, and drug or test naïve) prior to treatment or testing. (This information can often be tabulated). | Figure 2. |
| Numbers analysed | 15 | a. Report the number of animals in each group included in each analysis. Report absolute numbers (e.g. 10/20, not 50%<sup>2</sup>).<br>b. If any animals or data were not included in the analysis, explain why. | Figure 5 and Figure 6, each diamonds points relevant to one animal. |
| Outcomes and estimation | 16 | Report the results for each analysis carried out, with a measure of precision (e.g. standard error or confidence interval). | Figure 5, 6, confidence intervals. |
| Adverse events | 17 | a. Give details of all important adverse events in each experimental group.<br>b. Describe any modifications to the experimental protocols made to reduce adverse events. | Results section, adverse events on reproductive parameters. |
| **DISCUSSION** |  |  |  |
| Interpretation/<br>scientific<br>implications | 18 | a. Interpret the results, taking into account the study objectives and hypotheses, current theory and other relevant studies in the literature.<br>b. Comment on the study limitations including any potential sources of bias, any limitations of the animal model, and the imprecision associated with the results<sup>2</sup>.<br>c. Describe any implications of your experimental methods or findings for the replacement, refinement or reduction (the 3Rs) of the use of animals in research. | Discussion<br>section. |
| Generalisability/<br>translation | 19 | Comment on whether, and how, the findings of this study are likely to translate to other species or systems, including any relevance to human biology. | Conclusion<br>. |
| Funding | 20 | List all funding sources (including grant number) and the role of the funder(s) in the study. | Acknowled<br>gments. |

